# Analysis and Visualization of Sleep Stages based on Deep Neural Networks

**DOI:** 10.1101/2020.06.25.170464

**Authors:** Patrick Krauss, Claus Metzner, Nidhi Joshi, Holger Schulze, Maximilian Traxdorf, Andreas Maier, Achim Schilling

**Affiliations:** Neuroscience Lab, Experimental Otolaryngology, University Hospital Erlangen, Germany; Chair of Biophysics, Friedrich-Alexander University Erlangen-Nürnberg (FAU), Germany; Department of Otolaryngology, Head and Neck Surgery, University Hospital Erlangen, Germany; Chair of Machine Intelligence, Friedrich-Alexander University Erlangen-Nürnberg (FAU), Germany

**Keywords:** sleep stage scoring, hypnodensity graphs, multidimensional scaling (MDS), electroencephalography (EEG), artificial neural networks, deep learning, polysomnography (PSG), sleep cycle analysis

## Abstract

Automatic sleep stage scoring based on deep neural networks has come into focus of sleep researchers and physicians, as a reliable method able to objectively classify sleep stages, would save human resources and thus would simplify clinical routines. Due to novel open-source software libraries for Machine Learning in combination with enormous progress in hardware development in recent years a paradigm shift in the field of sleep research towards automatic diagnostics could be observed. We argue that modern Machine Learning techniques are not just a tool to perform automatic sleep stage classification but are also a creative approach to find hidden properties of sleep physiology. We have already developed and established algorithms to visualize and cluster EEG data, in a way so that we can already make first assessments on sleep health in terms of sleep-apnea and consequently daytime vigilance. In the following study, we further developed our method by the innovative approach to analyze cortical activity during sleep by computing vectorial cross-correlations of different EEG channels represented by hypnodensity graphs. We can show that this measure serves to estimate the period length of sleep cycles and thus can help to find disturbances due to pathological conditions.

## Introduction

Sleep stage scoring is a standard procedure and part of every polysomnographic analysis [1, 2]. Up to now, sleep stage scoring based on physiological signals (EEG: electroencephalography, EMG: electromyography, EOG: electroocculugraphy) is performed by experienced clinicians, which do the classification by hand according to the AASM guidelines [3]. However, this procedure is time consuming and highly prone to errors emphasized by high inter-rater variability [4–6]. To overcome these limitations Machine Learning algorithms were applied. These algorithms are still not fully trusted by clinicians as they act as a black-box, from which no one knows what the internal criteria for the different sleep stages really are. Thus, one core-problem of modern Machine Learning research has met clinical routines, the so called black-box problem [7] or depending on the scientific field the opacity debate [8]. This problem can be tackled by simply using hand-crafted features such as time tags of K complexes or sleep spindles as neural network input [9, 10], (for an example see [11]). However, this procedure is very elaborate and the results are still not satisfying. In times of rising computing power and increasing storage capacities, a Big Data approach with neural networks finding self organized the best features seems to be more promising. In order to analyze these features and emerging internal representations, novel concepts for data visualization are needed. A sophisticated dimensionality reduction method for visualising high dimensional data is a demanding task, as the certain popular methods can lead to pseudo clustering of data points caused by the initial conditions and the used hyper-parameters of the projection algorithm. A famous example for such phenomena is t-distributed stochastic neighbor embedding (t-SNE, [12]), which is highly unstable and extremely dependent on the initialization and choice of parameters [13].

In previous studies, we developed several approaches to statistically analyze and visualize high-dimensional neural data [14, 15]. We developed a statistical method for analyzing and comparing high-dimensional spatiotemporal cortical activation patterns for different auditory and somatosensory stimulus conditions in both, rodents and humans [14]. The cortical activity patters were represented by amplitude vectors calculated via a sliding window method (for the exact procedure see [14]). We could already demonstrate that this method enables to discriminate different sleep stages in human EEG recordings [16]. Furthermore, we could analyze the microstructure of cortical activity during sleep and found that it reflects respiratory events and state of daytime vigilance [17]. Recently, our method has been generalized, and can now be used to analyze and compare representations of artificial neural networks [15]. In the following study, we first illustrate how to visualize the representations of EEG data gained from different layers of artificial neural networks (sleep stage embeddings). Furthermore, we demonstrate that these complex representations cluster better in higher layers of the artificial neural networks, quantified by the generalized discrimination value (GDV, see also [15]). Subsequently, we visualize the output of the last layer of the neural network, and thus the momentary probabilities of the predicted sleep stages, as so called hypnodensity graphs (as introduced by [18]). Finally, we use these probability vectors to calculate vectorial cross-correlations in order to analyze the period length of sleep cycles.

## Methods

### Data base

The study was conducted in the Department of Otorhinolaryngology, Head Neck Surgery, of the Friedrich-Alexander University Erlangen-Nürnberg (FAU), following approval by the local Ethics Committee (323–16 Bc). All participants were recruited by the Department of Otorhinolaryngology, Head and Neck Surgery. Written informed consent was obtained from the participants before the cardiorespiratory polysomnography (PSG). Inclusion criterion for this study was age between 21 and 80 years. Exclusion criteria were a positive history of misuse of sedatives, alcohol or addictive drugs and untreated sleep disorders. Data analysis was carried out during “time in bed” of the subjects.

### Sleep stage classification with neural networks

A deep neural network was trained on single channel sleep EEG data. We used a total of 68 data sets, of which 54 were training data sets and 14 were test data sets. Specifically, we used a network consisting of several convolutional layers and two bidirectional stateful LSTM layers, which was trained with error back propagation to classify the sleep stages (for exact configuration see Figure 1).

**Figure 1:**
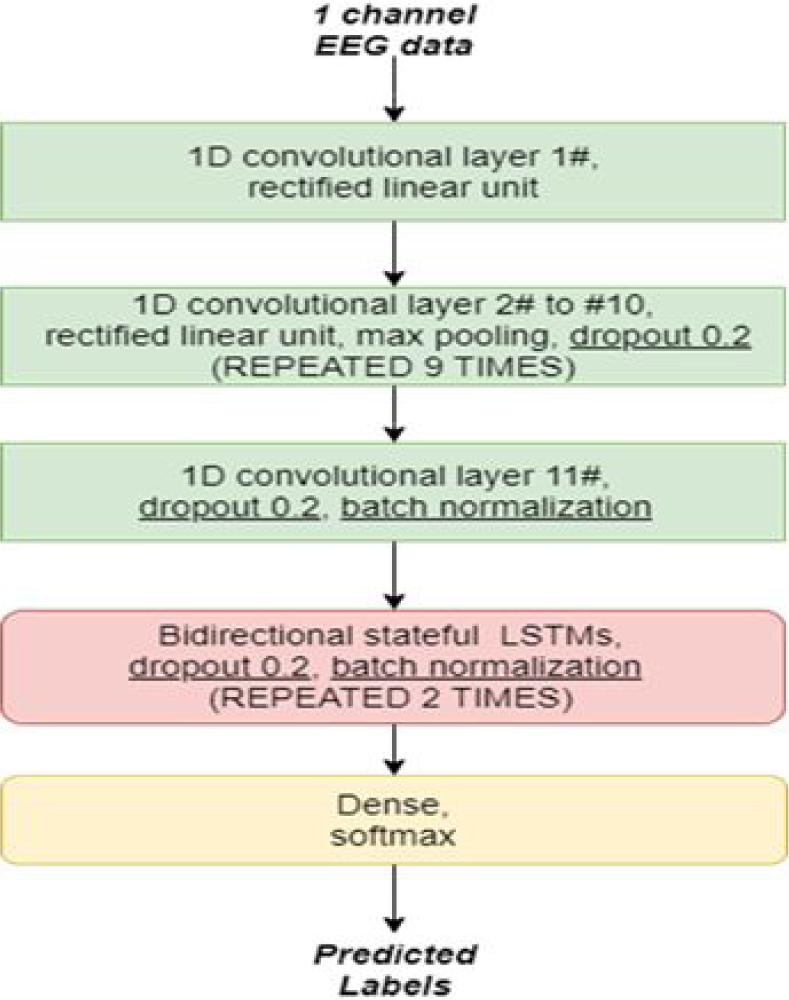
Network architecture. Building blocks of the deep neural network trained on sleep stage classification of EEG data. The networks consists of eleven 1D convolutional layer, nine max pooling layers, 2 layers of bidirectional stateful LSTMs, and a fully connected classification layer with softmax output.

### Software resources

The software is written using the programming language python 3.6, together with the KERAS AI library [19] and Tensorflow backend. The data is pre-processed using Numpy [20]. For multidimension-scaling (MDS) the scikit-learn library [21] is used and the visualization of the results is done using the matplotlib library [22].

### Hypnodensity correlations

The hypnodensity cross-correlation for lag-time *τ* is defined as

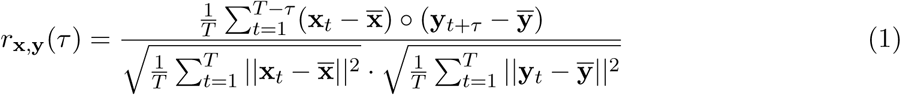

and the hypnodensity auto-correlation for lag-time *τ* is defined as

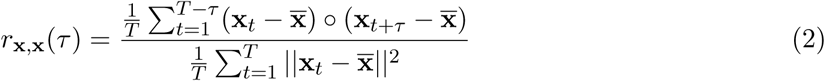

with 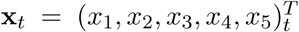 and 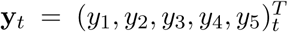 being 5-dimensional vectors that contain the momentary probabilities of all sleep stages (indices 1 to 5) at time *t* for EEG channels *x* and *y*, respectively, and the mean probability vectors 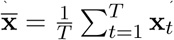 and 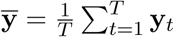. Here, ∘ is the dot product, and *‖ ‖* the vector norm.

## Results

### Sleep stage embeddings

We could show that even a single EEG channel contains enough information to classify different sleep stages. We used several convolutional blocks, which transform the input EEG data in high dimensional complex features. Thus, in contrast to handcrafted features we make the neural network to find its own features. This higher order features lead to a good separability of the transformed EEG vectors (neural network layer output or representation) belonging to different sleep stages. In the following, we refer to this using the term *sleep stage embeddings*.

We visualize the sleep stage embeddings by a dimensionality reduction into 2D using multidimensional scaling (MDS) [14] (Figure 2) and evaluate the generalized discrimination value (GDV) which quantifies separability of data classes in high-dimensional state spaces [15]. The MDS plots show that the convolutional layer lead to better separability (Figure 2 b-g) compared to z-scored raw EEG data (network input, Figure 2 a). Furthermore, the LSTM layer transforms the data so that the different classes are linearly separable (Figure 2 h). The softmax classification layer norms the data (value range [0,1]). The shape of the clusters in Figure 2 i can be explained by the fact that the softmax layers places the different classes at the 5 edges of a 4D-hyper-plane (example for 3D projections shown in Figures 3 and 4). As the probabilities of all 5 sleep stages have to sum up to a value of 1, the range of possible softmax outputs is located on this hyper-plane. The MDS projection of the hyper-plane in 2D leads to the shown patterns (Figure 2 i).

**Figure 2:**
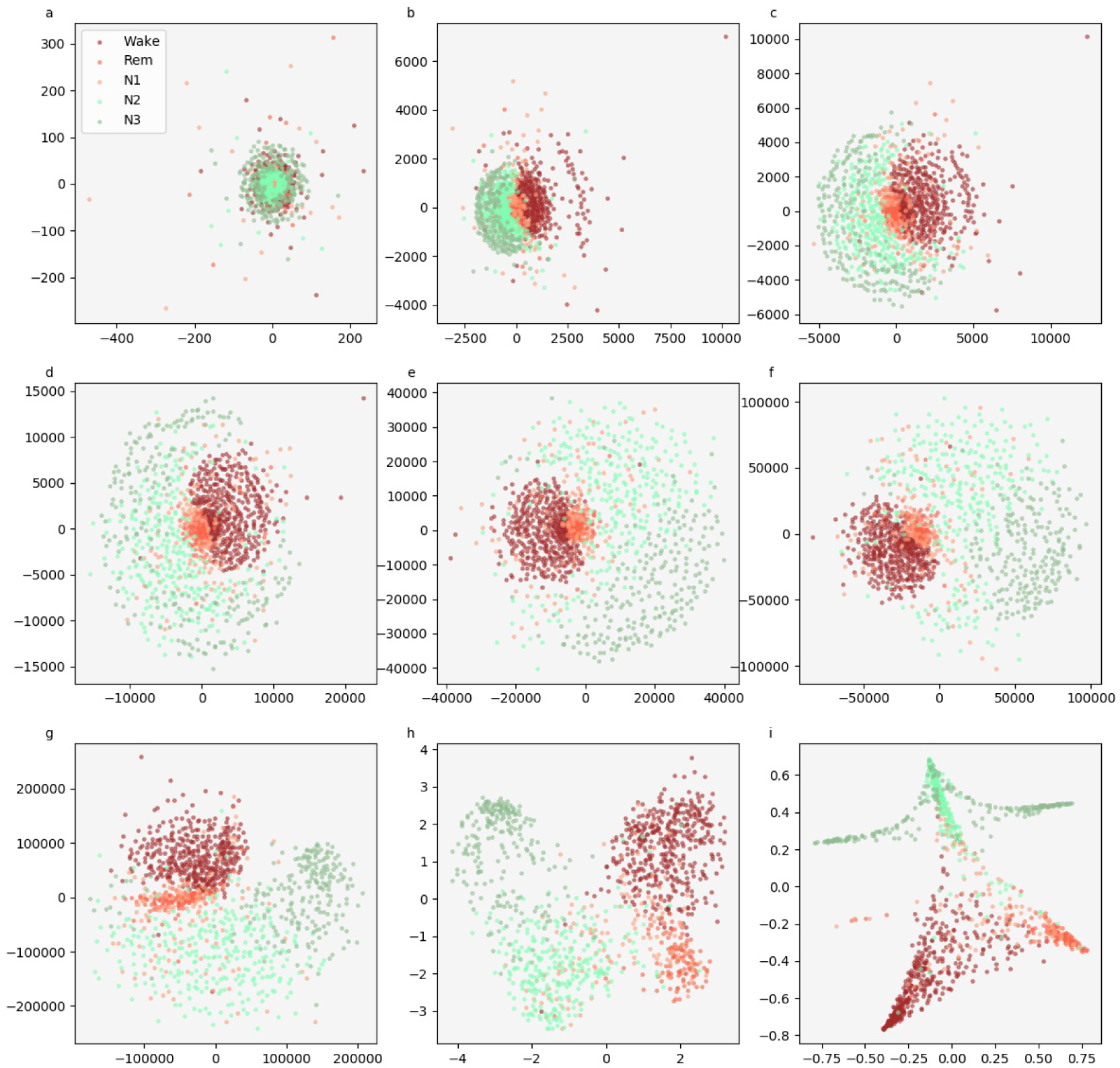
Sleep stage embeddings. MDS visualization of EEG data representations within the hidden layers of the deep neural network: input layer (a), max pooling layers (b-g), last LSTM layer (h), and softmax layer (i).

**Figure 3:**
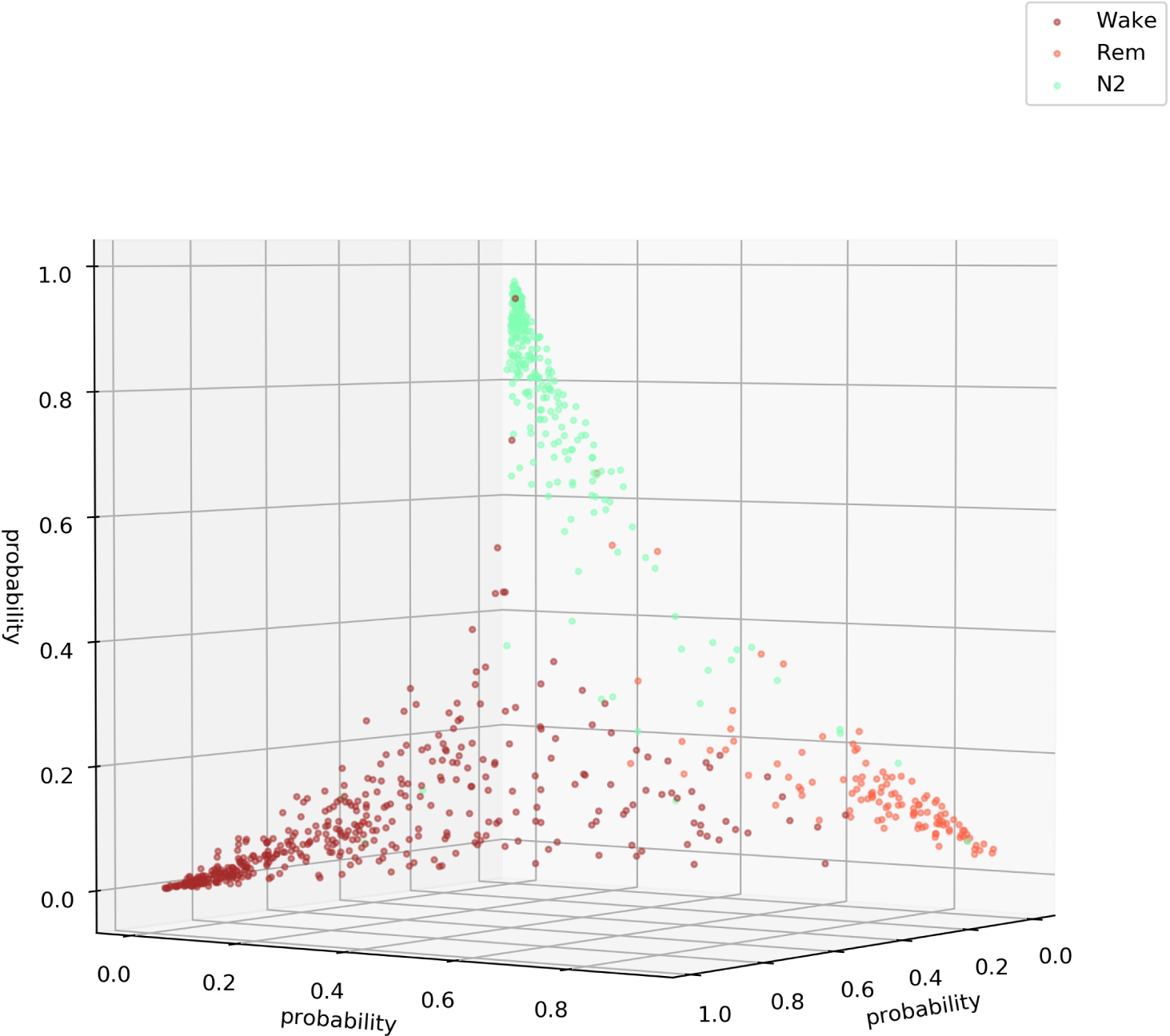
Softmax output for three sleep stages (Wake, REM, N2) This figure illustrates that the softmax output spans a hyper-plane (in 3D a 2D hyper-plane with 3 corners). Note that the possible outputs lie in the volume below the hyper-plane. Not all points lie on the plane as confusions with sleep stage N3 or N1 cannot be shown in a 3D plot. Thus, if the softmax output would for example point to sleep stage N3 the point would lie near the origin (0,0,0).

**Figure 4:**
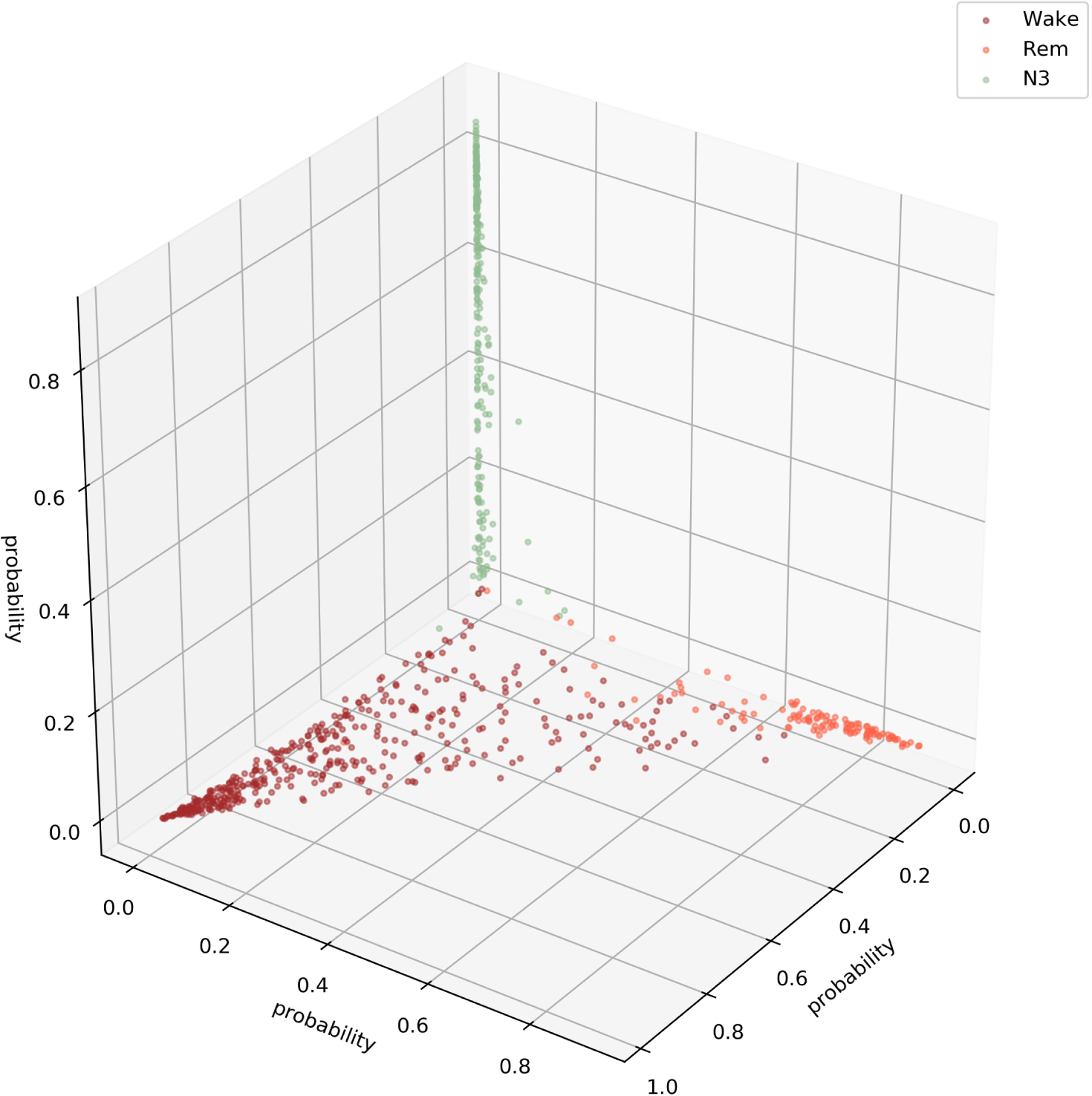
Softmax output of for three sleep stages (Wake, REM, N3) This figure illustrates that the softmax output spans a hyper-plane (in 3D a 2D hyper-plane with 3 corners). It can be seen that the sleep stage N3 is not confused with the Wake or the REM state.

The quantification of the class separability via the GDV (Figure 2, red dots: input, 6 maxpooling layers, the second LSTM layer, the softmax output layer), shows that the convolutional layers perform data pre-processing. In a previous study, we could show that the GDV has to overcome an “energy barrier”, when the data structure is complex. Thus, the number of layers where the GDV only slightly decreases depends on data complexity. As complex features are needed to classify the data, the number of layers with relatively small GDV decrease is high (cf. Figure 5). The LSMT layer leads to a clear increase of separability, i.e. drop in the GDV value (8th red point in Figure 5, compare also Figure 2 i). The softmax layer causes a further decrease of the GDV value.

**Figure 5:**
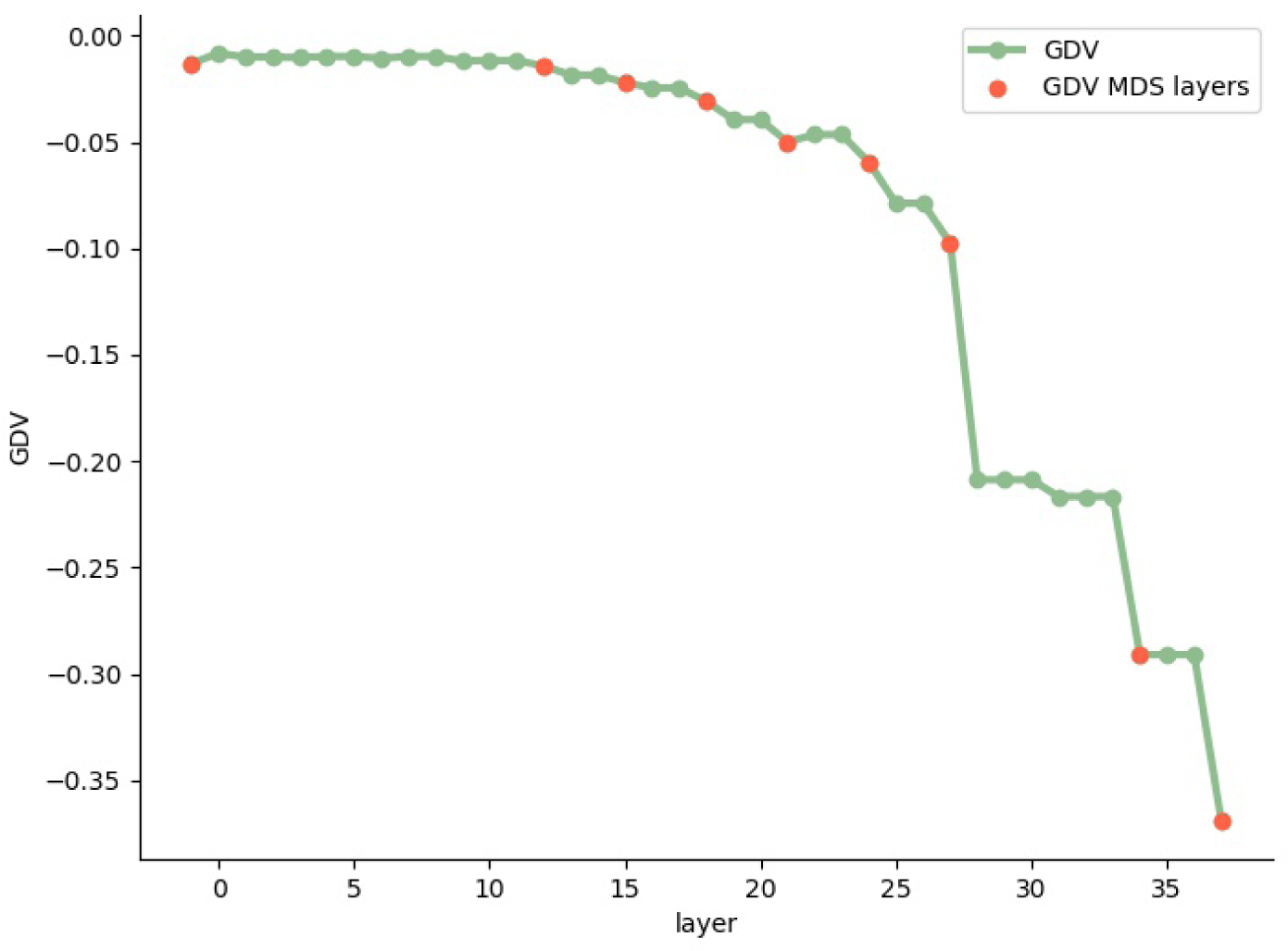
Separability of sleep stages. Separability increases with increasing layer depth. Note that, a GDV of 0 corresponds to non-separable data classes, whereas a GDV of −1 corresponds to a perfect data class separation. Red dots refer to the layer outputs shown in 2 (first red dot: input, 2nd-7th: higher max pooling layers, 8th: 2nd LSTM layer, 9th: softmax layer).

### Hypnodensity graphs

An efficient illustration of the softmax layer output are so called hypnodensity graphs [18]. Here, the probabilities for all sleep stages at each time point are shown instead of only the most probable sleep stage as it is done in traditional hypnograms. This is an elegant approach, since the probabilities (output of softmax layer) can be compared with the uncertainties, i.e. different assessments of different somnologists, which cause the relatively high inter-rater-variability [4–6].

Additionally, the automatic sleep stage classification can be used to evaluate the data with higher temporal resolution. Even though, the network has been trained on labeled data with a temporal resolution of 30 seconds which is in accordance with AASM scoring rules [3], we evaluate sleep stage probabilities predicted by the neural network with a temporal resolution of 5 seconds using a sliding window approach (Figures 6 and S1 – S10).

**Figure 6:**
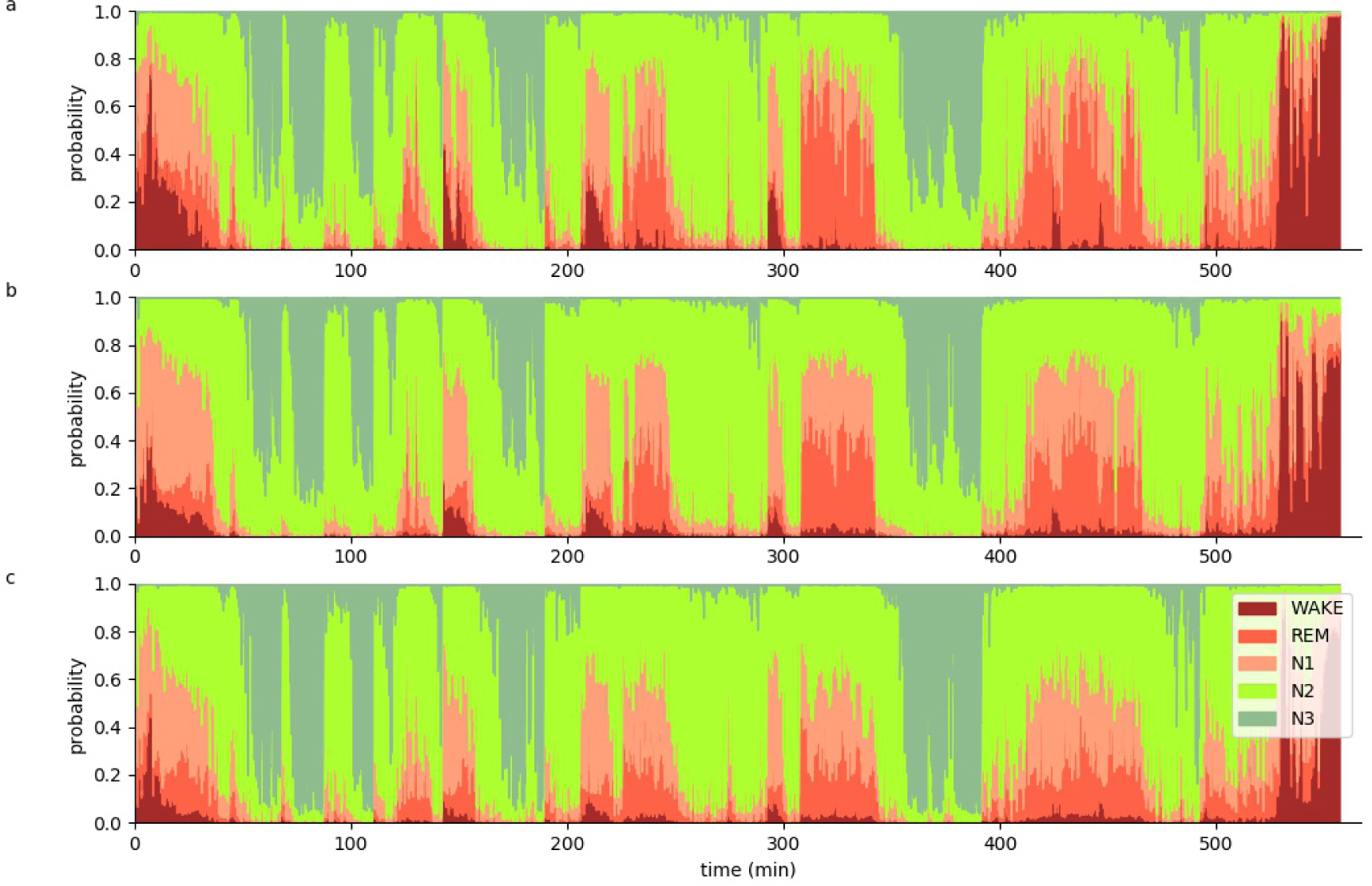
Hypnodensity graph. Hypnodensity graph of subject 55 with a temporal resolution of 5 seconds separately evaluated for the three different EEG channels C4 (a), F4 (b) and O2 (c).

The hypnodensity plots show that each EEG channel is sufficient to perform sleep stage classification, as the hypnodensity graphs are similar (Figure 6 a-c) for all the channels. This fact can be further emphasized by the calculation of pairwise cross-correlations as follows.

### Hypnodensity based sleep cycle period length analysis

On the basis of these EEG channel specific hypnodensity graphs, sleep stage probabilities across and within channels can be compared (see Methods). For this purpose we calculate generalized vector auto- and cross-correlations of the 5-dimensional probability vectors for the 3 EEG channels of each data set. This correlations provide a lot of information on neuronal activity patterns during sleep. Thus, all auto-as well as cross-correlations have a global maximum, which is near to the value 1, at 0 lag time. This means that there is no time shift of the neuronal patterns between the different channels. Furthermore, the high correlation coefficients at lag time 0 indicate that all channels are sufficient to perform sleep scoring, which could already been predicted by the hypnodensity graphs. Local side-maxima indicate repetitions of neuronal patterns as they occur in repeating sleep-cycles.

For subject 55, the auto and cross-correlations show a local maximum at a lag-time of about 100 minutes (Figure 7). This means that after this period of time the sleep stages repeat. In contrast, data of subject 59 show two local maxima at lag-times of 75 and 150 minutes, respectively (Figure 8). Hence, this subject’s sleep stage period length is about 75 minutes. This means that our novel method enables estimating the individual period length of sleep cycles. Further data are shown in Figures S11 – S19.

**Figure 7:**
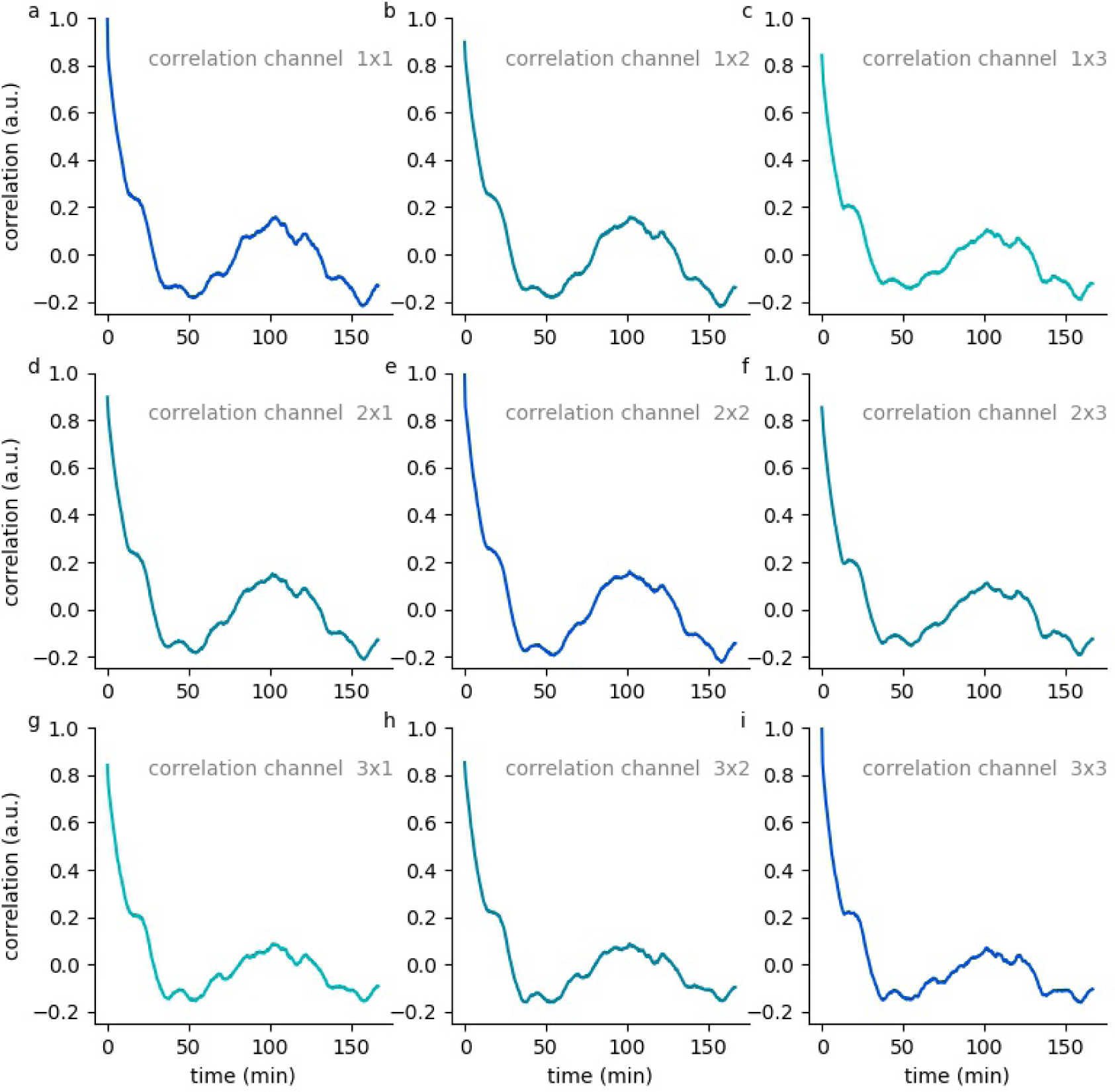
Hypnodensity cycles. Temporal auto- and cross correlations of 5-dimensional hypnodensity probability vectors of sleep stages. In subject 55, a local maximum at a lag-time of about 100 minutes can be observed, indicating an individual period length of sleep cycles of 100 minutes.

**Figure 8:**
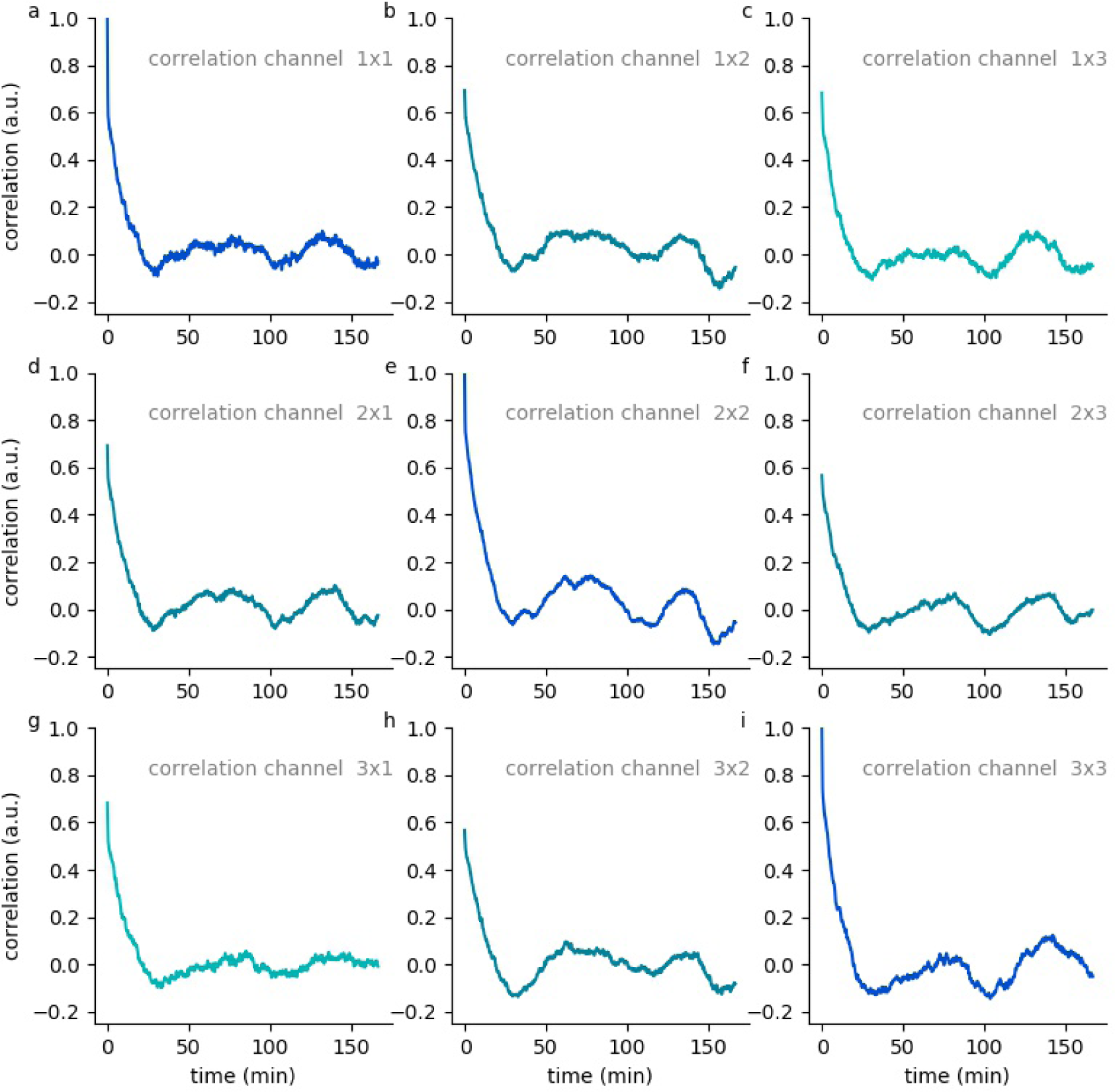
Hypnodensity cycles. Temporal auto- and cross correlations of 5-dimensional hypnodensity probability vectors of sleep stages. In subject 59, two local maxima at lag-times of about 75 and 150 minutes can be observed, indicating an individual period length of sleep cycles of 75 minutes.

## Discussion

In this study, we present novel approaches for the evaluation and visualization of neural data recorded during human sleep. In contrast to numerous previous studies and network architectures used to perform automatic sleep stage scoring (e.g. [23–28]), we provide approaches helping to interpret and understand the network output.

Thus, we illustrate how deep neuronal networks automatically find features that help to separate different sleep stages based on single channel EEG data. That this representations actually lead to a good separability of the different sleep stages is visualized using MDS plots and quantified using the GDV value.

The softmax output is visualized in hypnodensity plots [18] a sophisticated way to show the uncertainties in sleep stage classification. The calculation of the vectorial correlations is a tool to gain insight into sleep cycles. Thus, we propose the correlation plots as novel tool to find disturbances in the period length of sleep cycles, and thus as an indicator for pathological conditions.

This study is in line with numerous other studies trying to use artificial neural networks as a model [29–32], and as a tool [33] to analyze and understand brain activity. This combined approach at the intersection of neuroscience and artificial intelligence is crucial to make further progress in the field, as traditional popular analysis methods have been proven to be not sufficient to understand brain dynamics [34–36].

## Additional Information

## Acknowledgments

This work was funded by the Deutsche Forschungsgemeinschaft (DFG, German Research Foundation): grant KR5148/2-1 to PK – project number 436456810, the Interdisciplinary Center for Clinical Research (IZKF) at the University Hospital of the University Erlangen-Nuremberg (grant ELAN-17-12-27-1-Schilling to AS), and the Emergent Talents Initiative (ETI) of the University Erlangen-Nuremberg (grant 2019/2-Phil-01 to PK). We thank NVidia for the donation of GPU devices for Machine Learning purposes.

## Supplements

**Figure S1:**
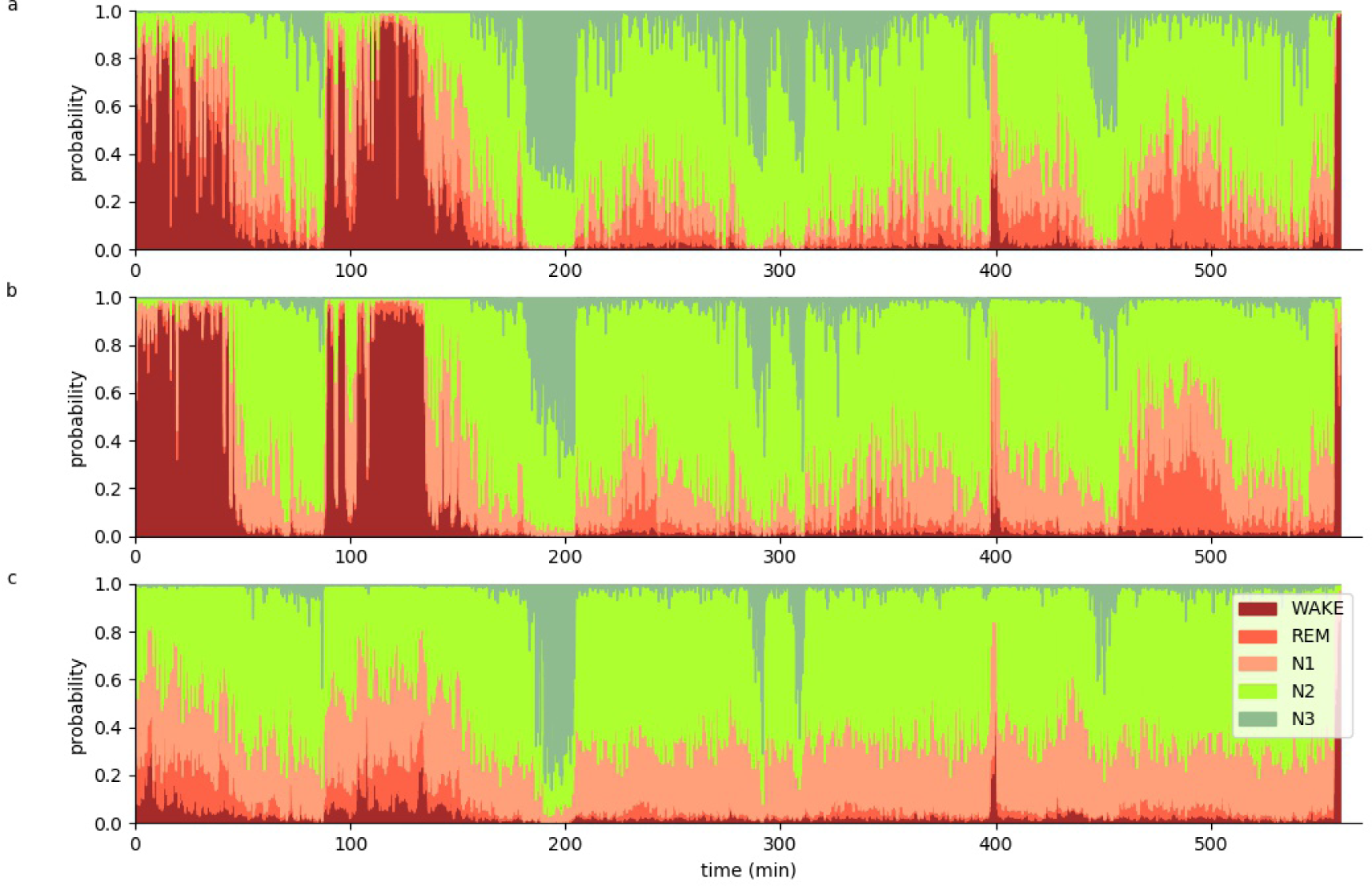
Hypnodensity graph. Hypnodensity graph of subject 56 with a temporal resolution of 5 seconds separately evaluated for the three different EEG channels C4 (a), F4 (b) and O2 (c).

**Figure S2:**
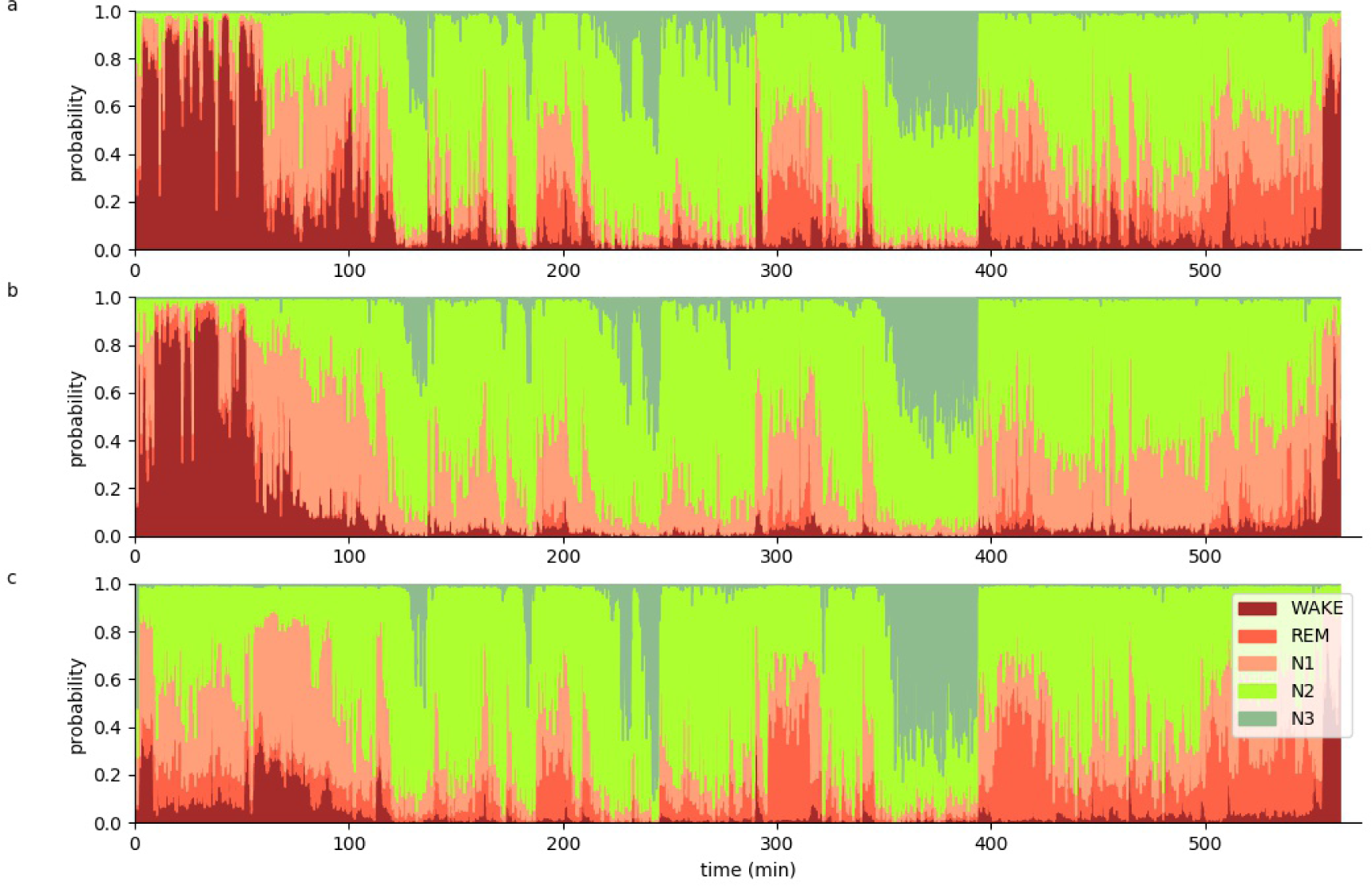
Hypnodensity graph. Hypnodensity graph of subject 57 with a temporal resolution of 5 seconds separately evaluated for the three different EEG channels C4 (a), F4 (b) and O2 (c).

**Figure S3:**
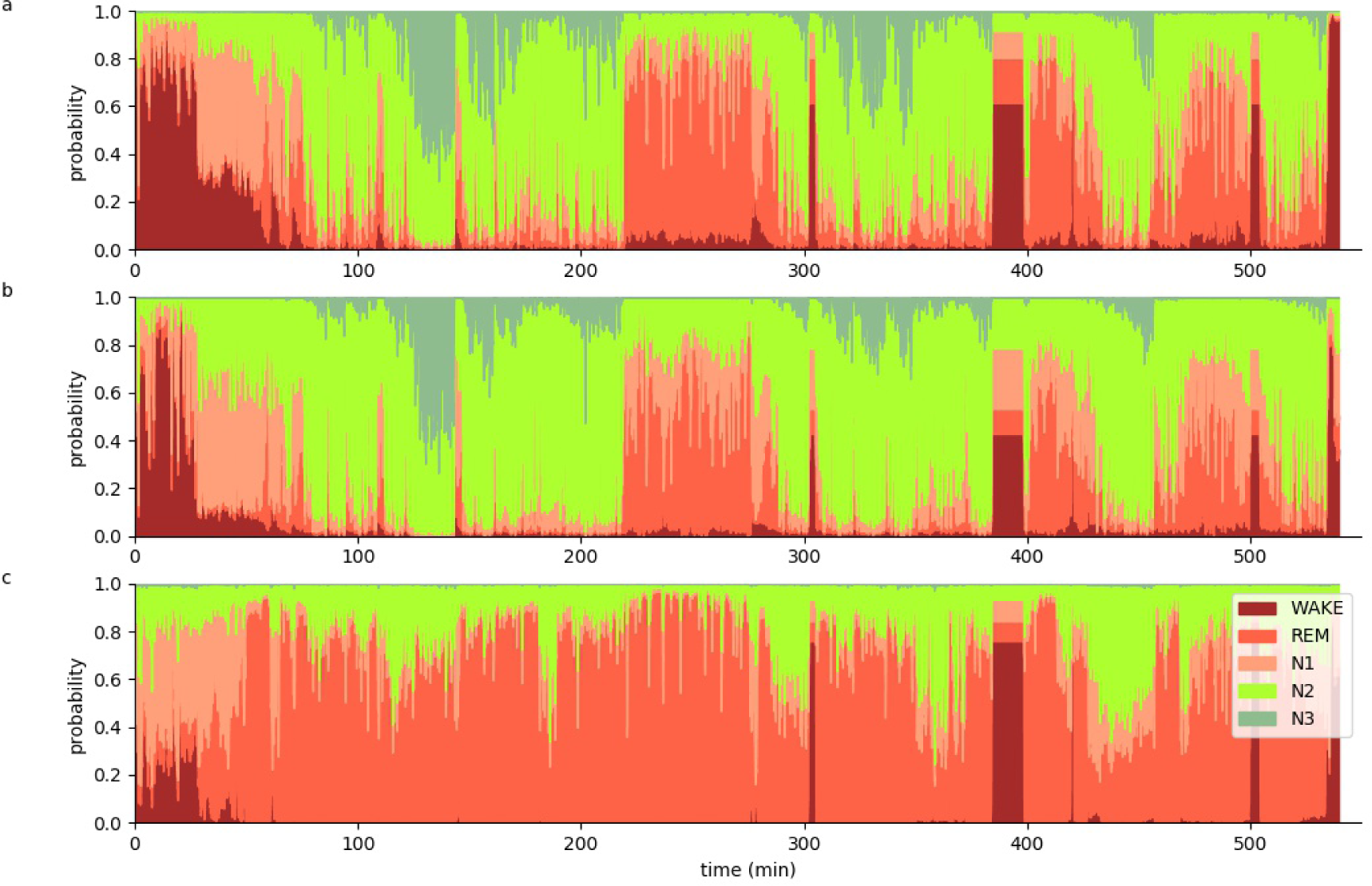
Hypnodensity graph. Hypnodensity graph of subject 58 with a temporal resolution of 5 seconds separately evaluated for the three different EEG channels C4 (a), F4 (b) and O2 (c).

**Figure S4:**
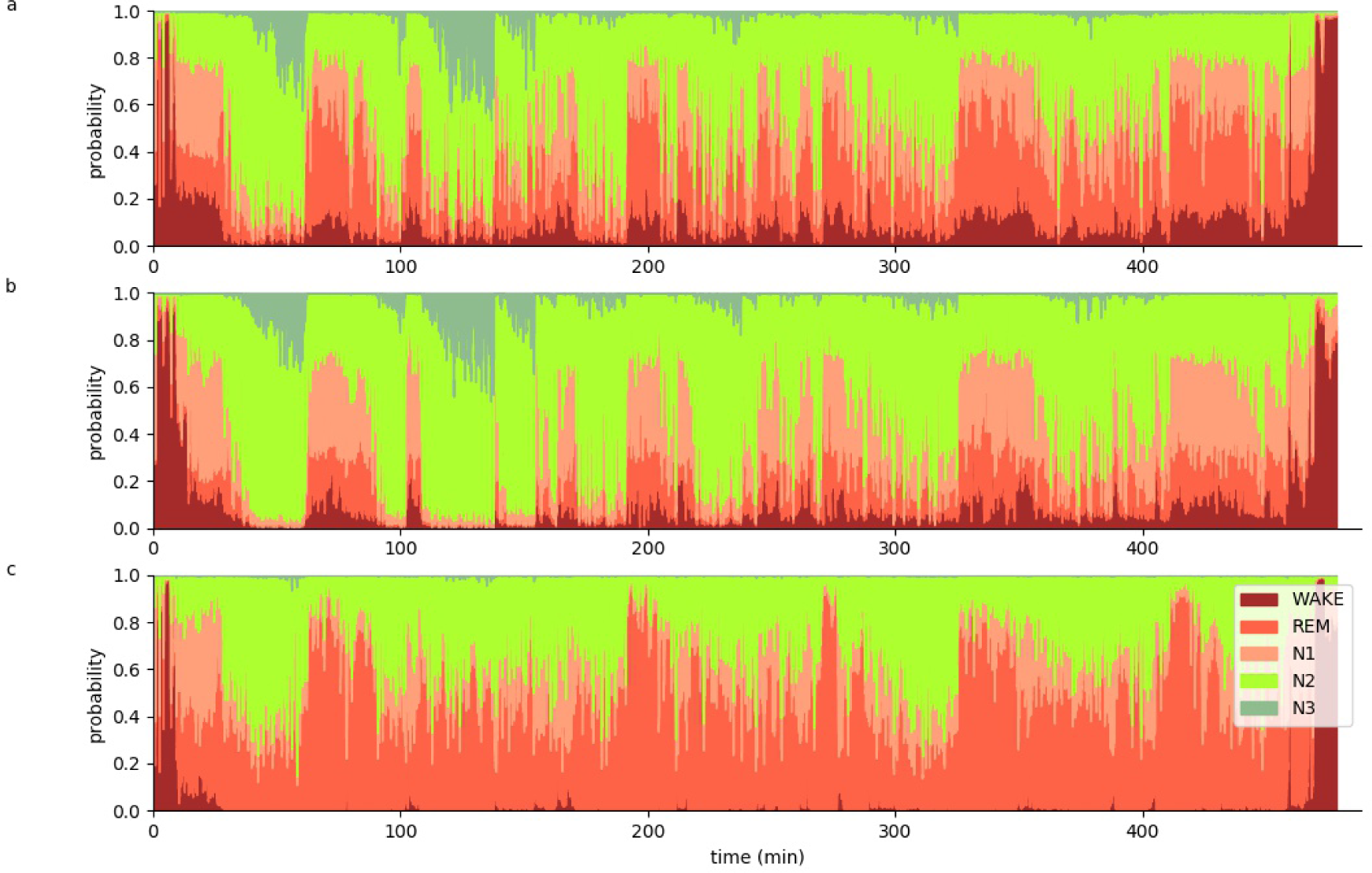
Hypnodensity graph. Hypnodensity graph of subject 59 with a temporal resolution of 5 seconds separately evaluated for the three different EEG channels C4 (a), F4 (b) and O2 (c).

**Figure S5:**
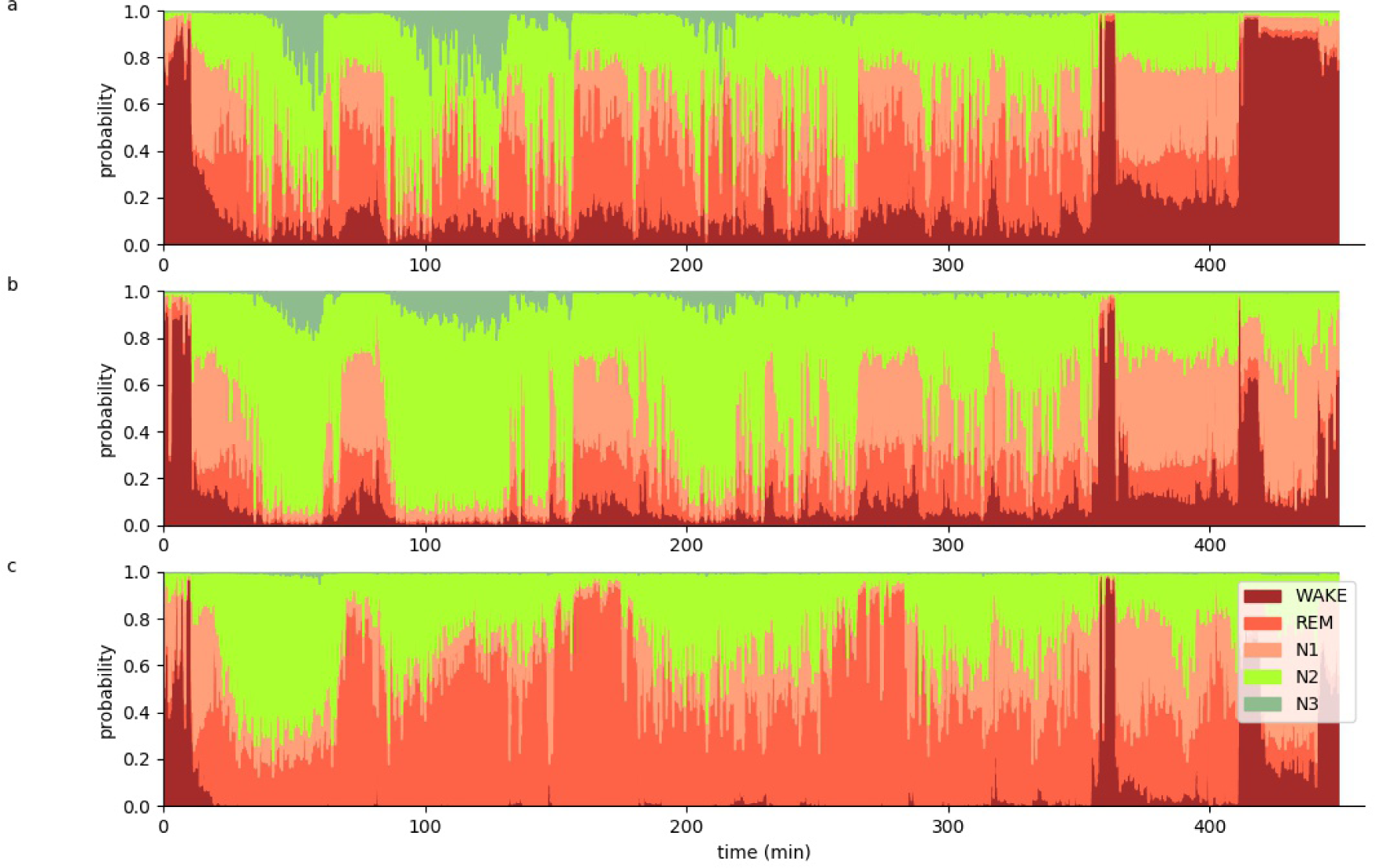
Hypnodensity graph. Hypnodensity graph of subject 60 with a temporal resolution of 5 seconds separately evaluated for the three different EEG channels C4 (a), F4 (b) and O2 (c).

**Figure S6:**
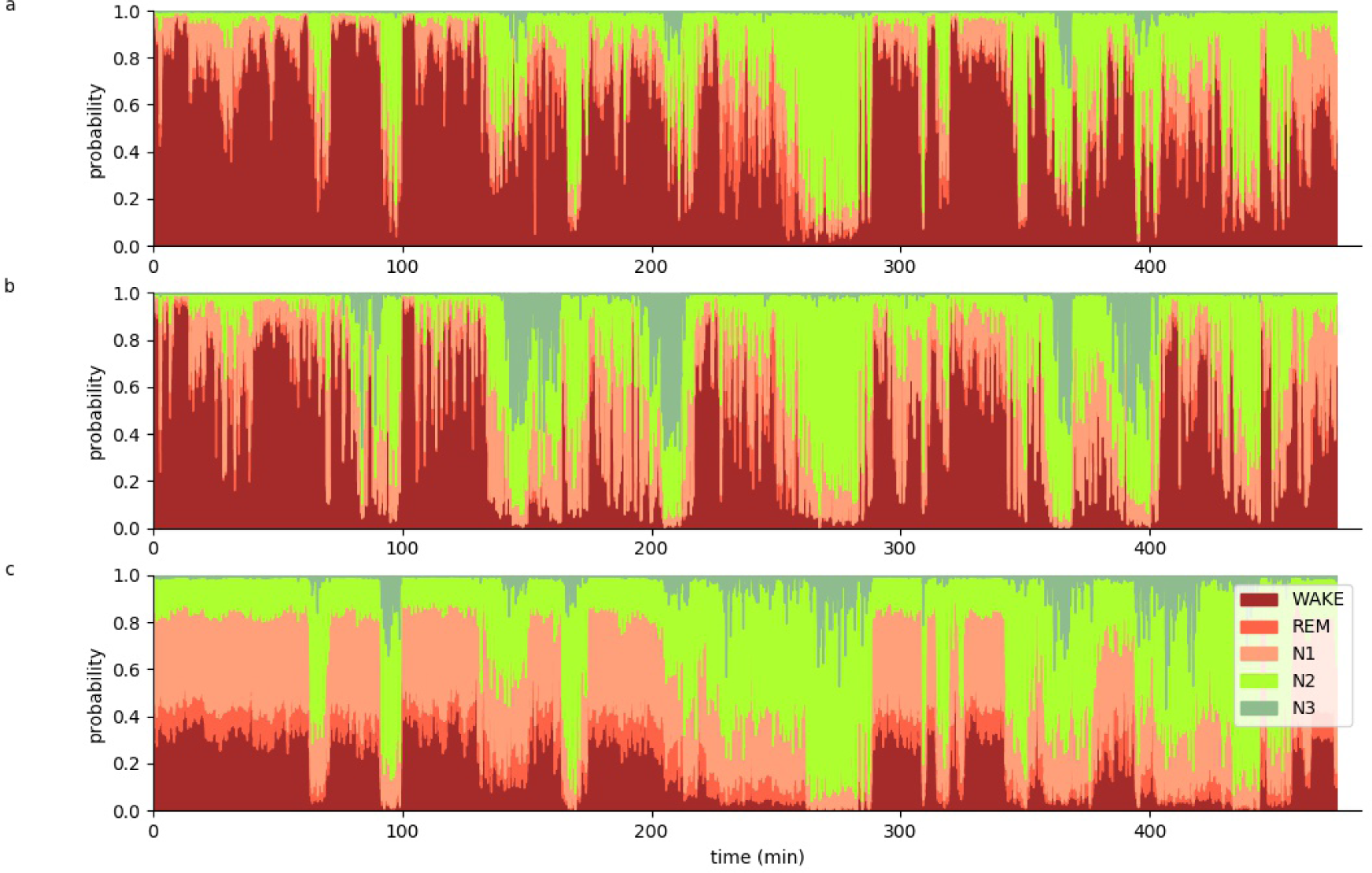
Hypnodensity graph. Hypnodensity graph of subject 61 with a temporal resolution of 5 seconds separately evaluated for the three different EEG channels C4 (a), F4 (b) and O2 (c).

**Figure S7:**
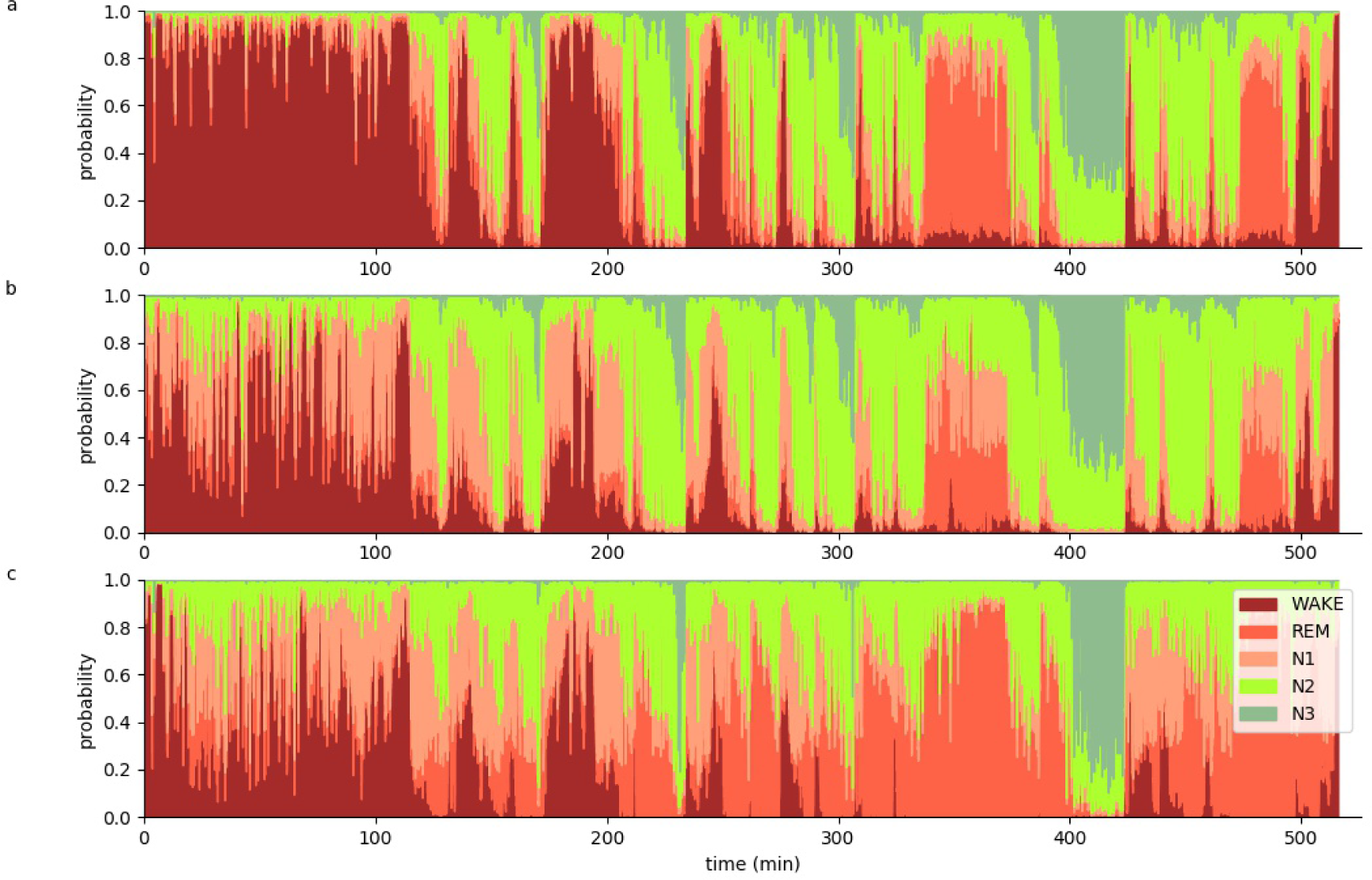
Hypnodensity graph. Hypnodensity graph of subject 62 with a temporal resolution of 5 seconds separately evaluated for the three different EEG channels C4 (a), F4 (b) and O2 (c).

**Figure S8:**
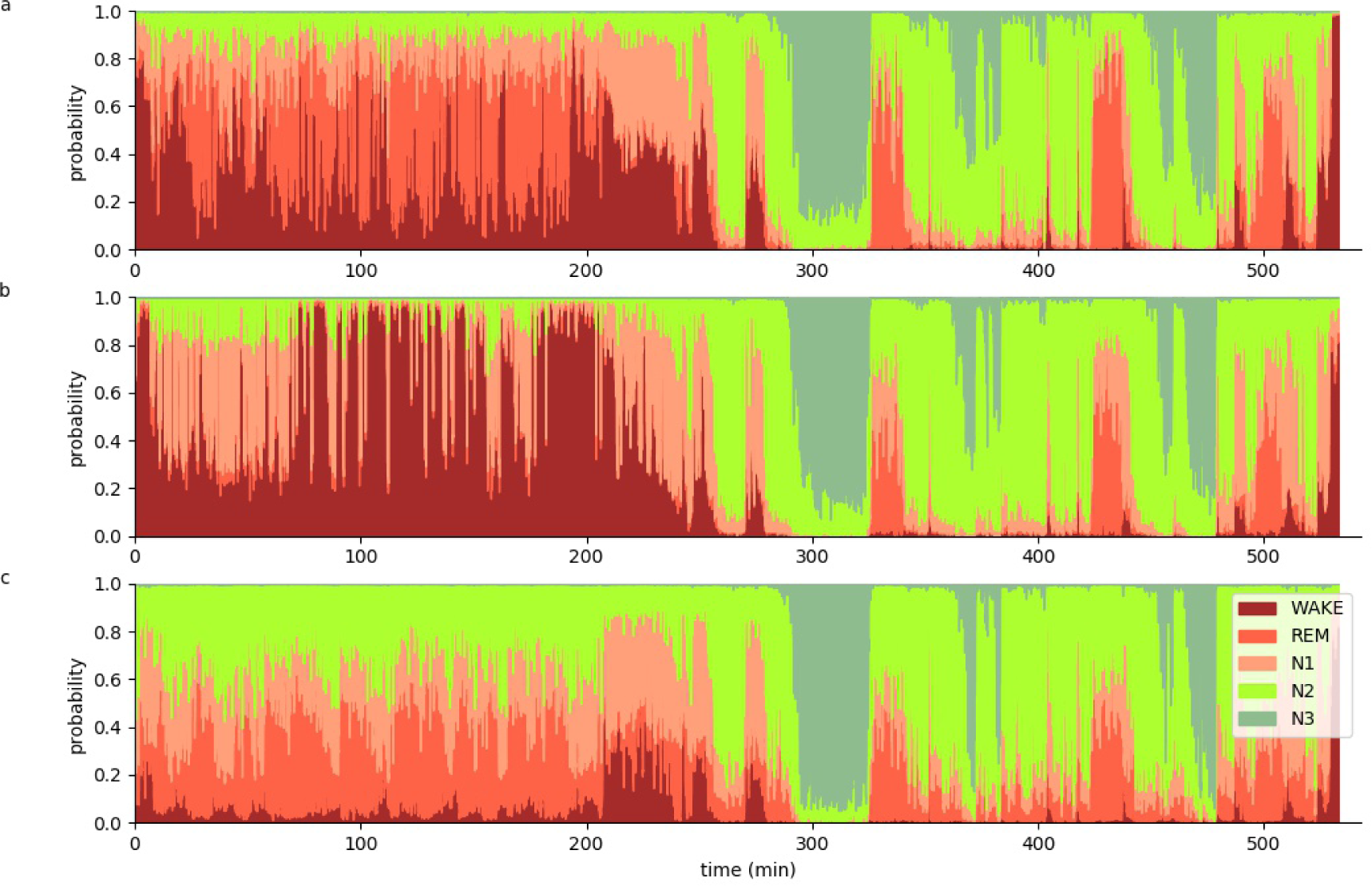
Hypnodensity graph. Hypnodensity graph of subject 63 with a temporal resolution of 5 seconds separately evaluated for the three different EEG channels C4 (a), F4 (b) and O2 (c).

**Figure S9:**
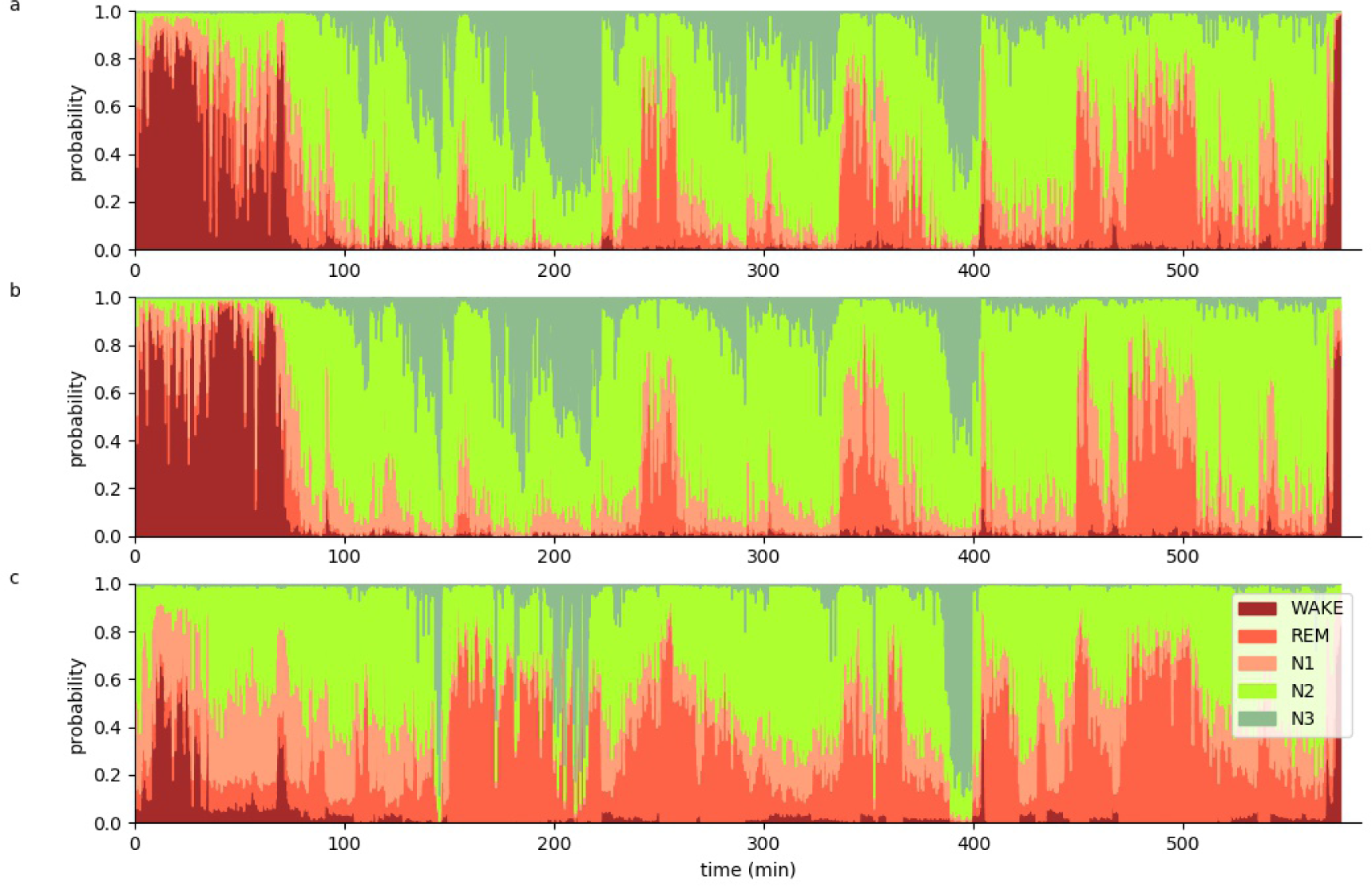
Hypnodensity graph. Hypnodensity graph of subject 64 with a temporal resolution of 5 seconds separately evaluated for the three different EEG channels C4 (a), F4 (b) and O2 (c).

**Figure S10:**
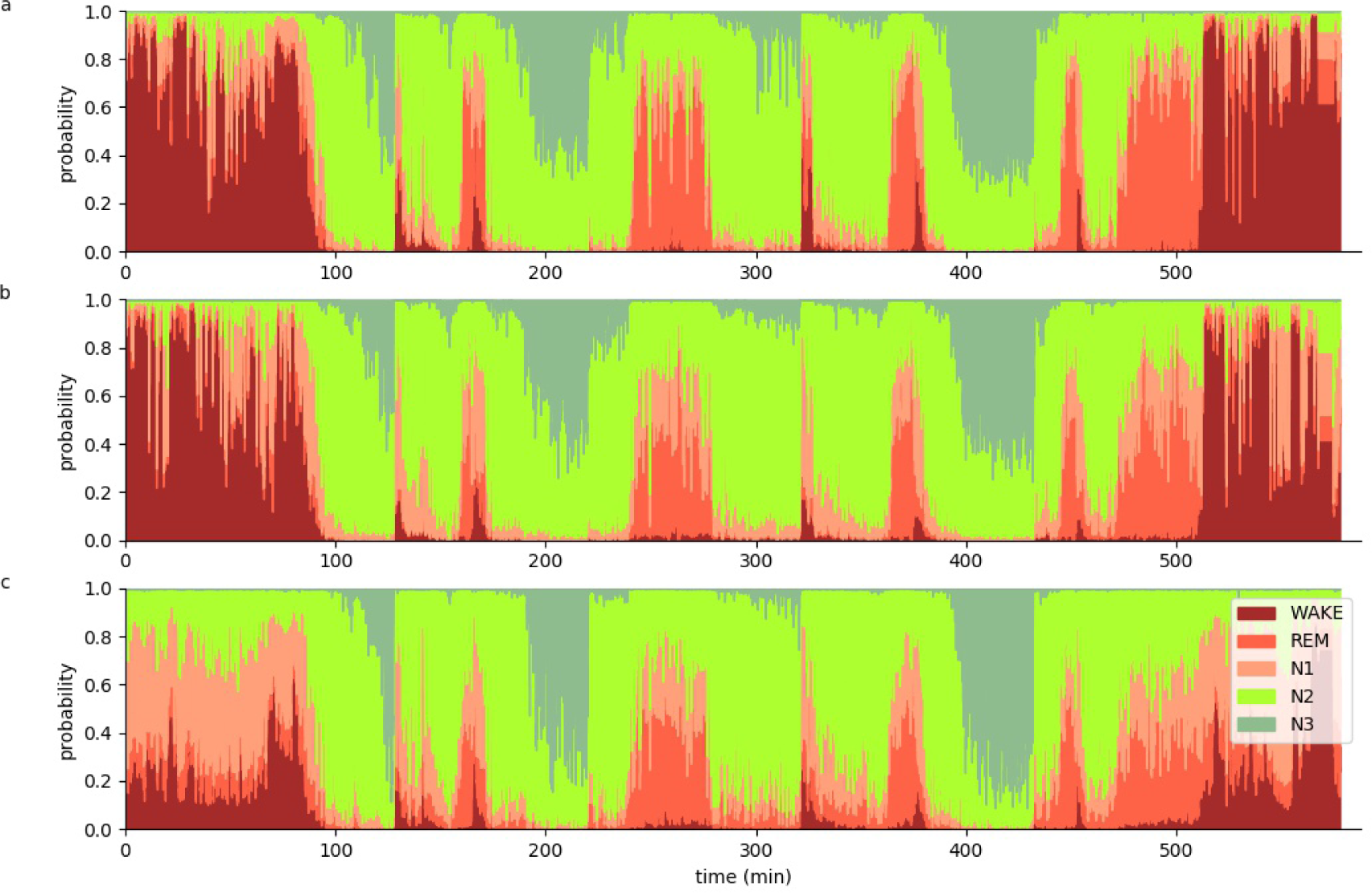
Hypnodensity graph. Hypnodensity graph of subject 65 with a temporal resolution of 5 seconds separately evaluated for the three different EEG channels C4 (a), F4 (b) and O2 (c).

**Figure S11:**
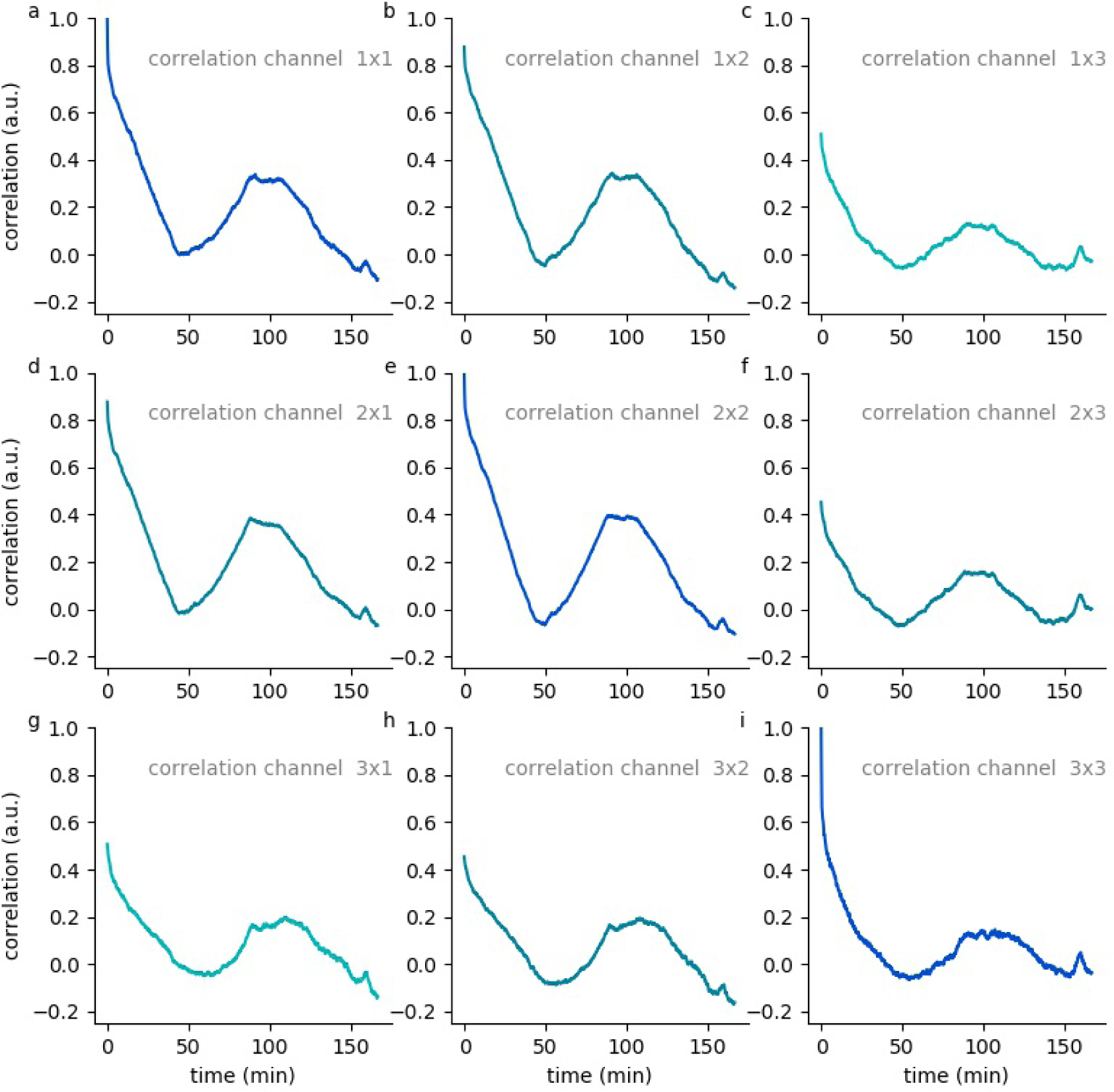
Hypnodensity cycles. Temporal auto- and cross correlations of 5-dimensional hypnodensity probability vectors of sleep stages. Shown are date from subject 56.

**Figure S12:**
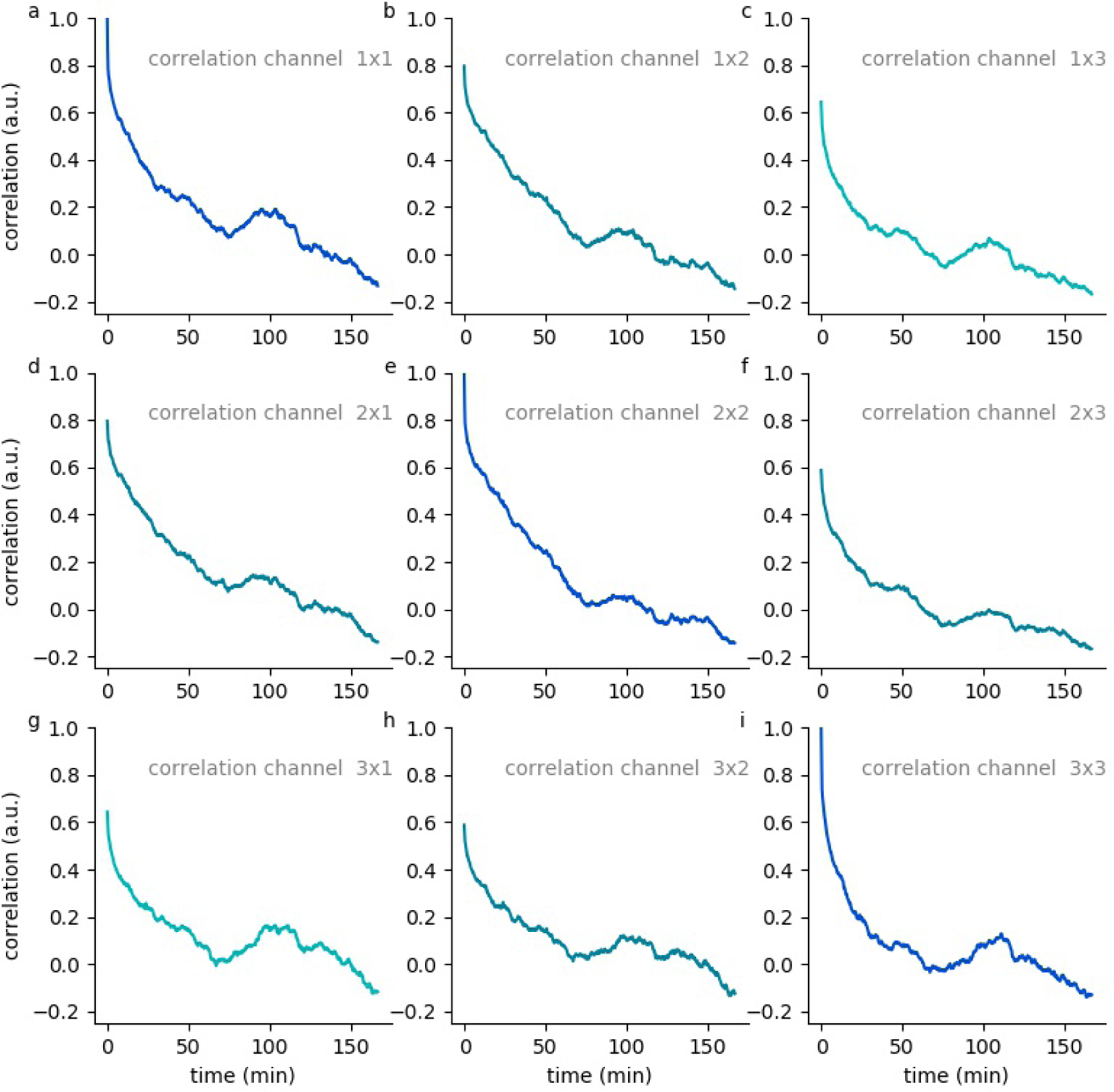
Hypnodensity cycles. Temporal auto- and cross correlations of 5-dimensional hypnodensity probability vectors of sleep stages. Shown are date from subject 57.

**Figure S13:**
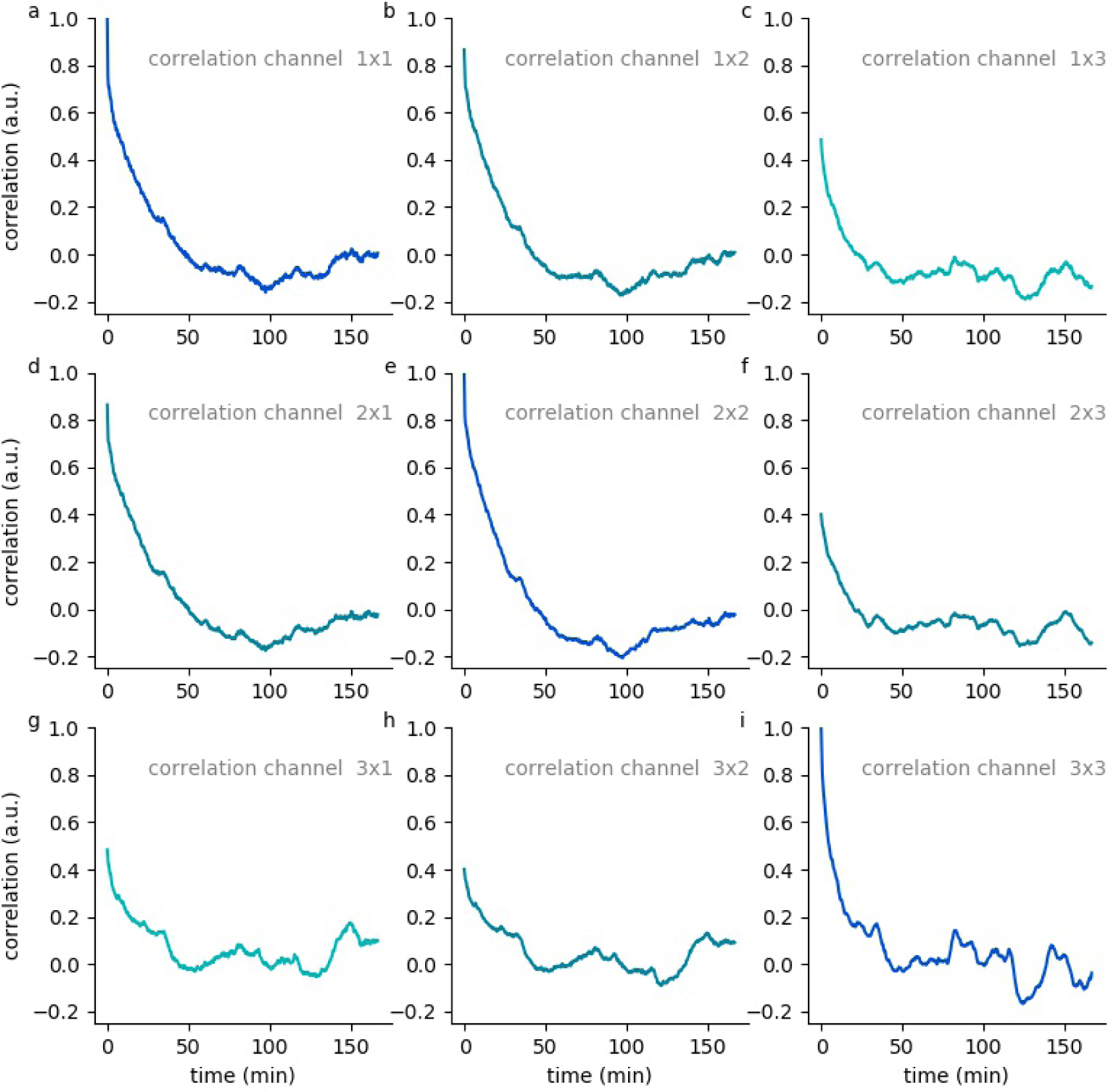
Hypnodensity cycles. Temporal auto- and cross correlations of 5-dimensional hypnodensity probability vectors of sleep stages. Shown are date from subject 58.

**Figure S14:**
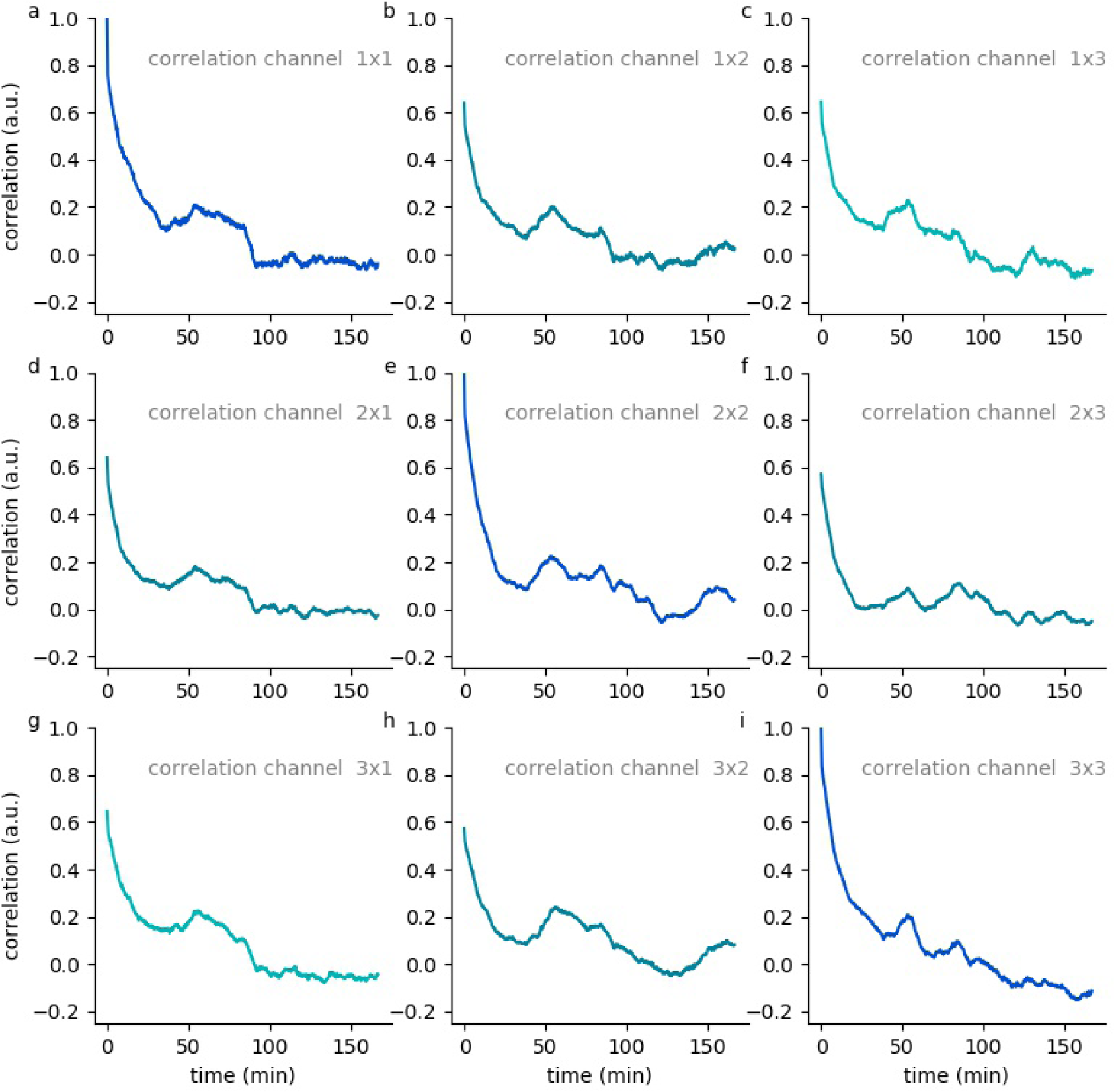
Hypnodensity cycles. Temporal auto- and cross correlations of 5-dimensional hypnodensity probability vectors of sleep stages. Shown are date from subject 60.

**Figure S15:**
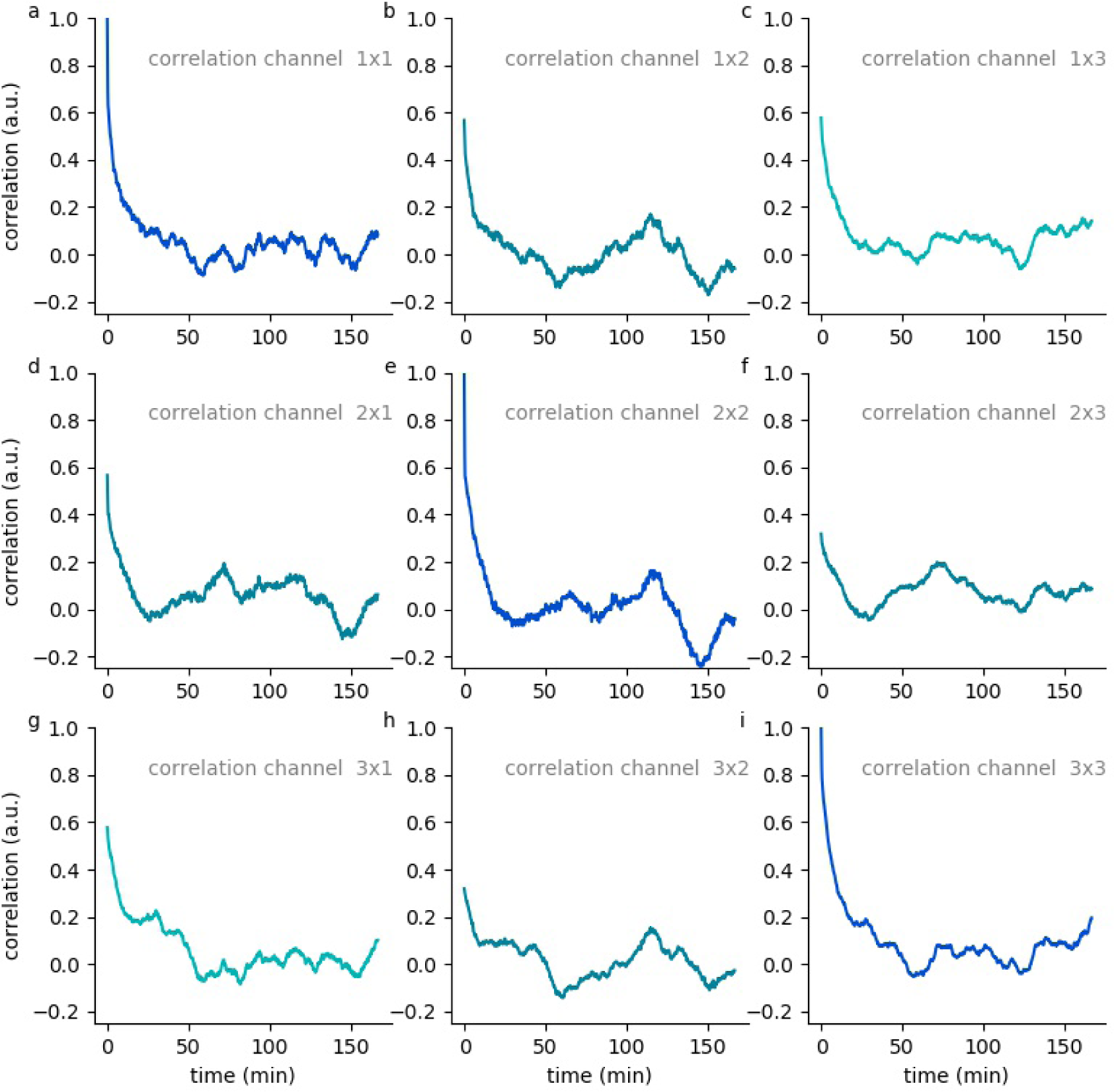
Hypnodensity cycles. Temporal auto- and cross correlations of 5-dimensional hypnodensity probability vectors of sleep stages. Shown are date from subject 61.

**Figure S16:**
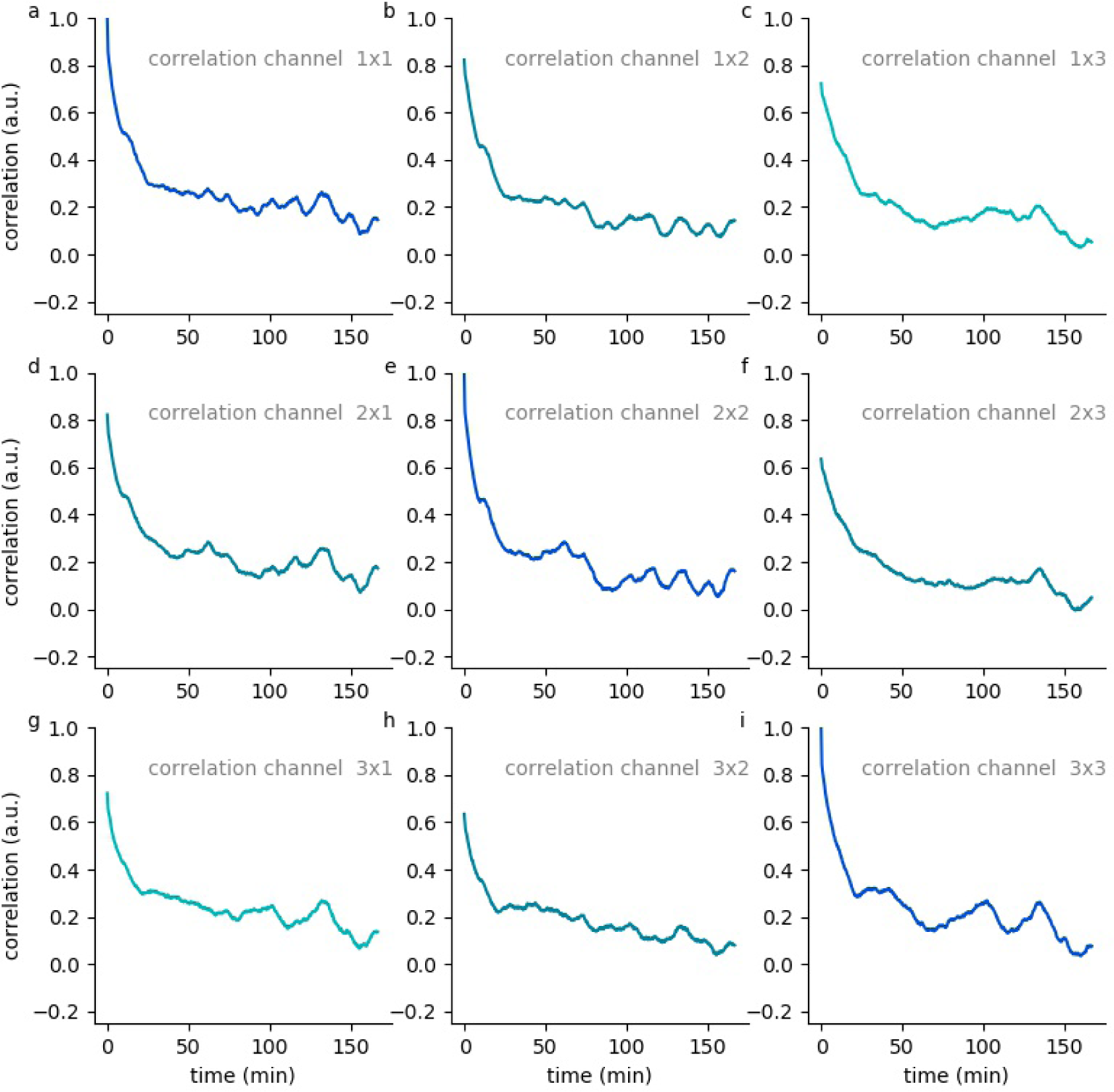
Hypnodensity cycles. Temporal auto- and cross correlations of 5-dimensional hypnodensity probability vectors of sleep stages. Shown are date from subject 62.

**Figure S17:**
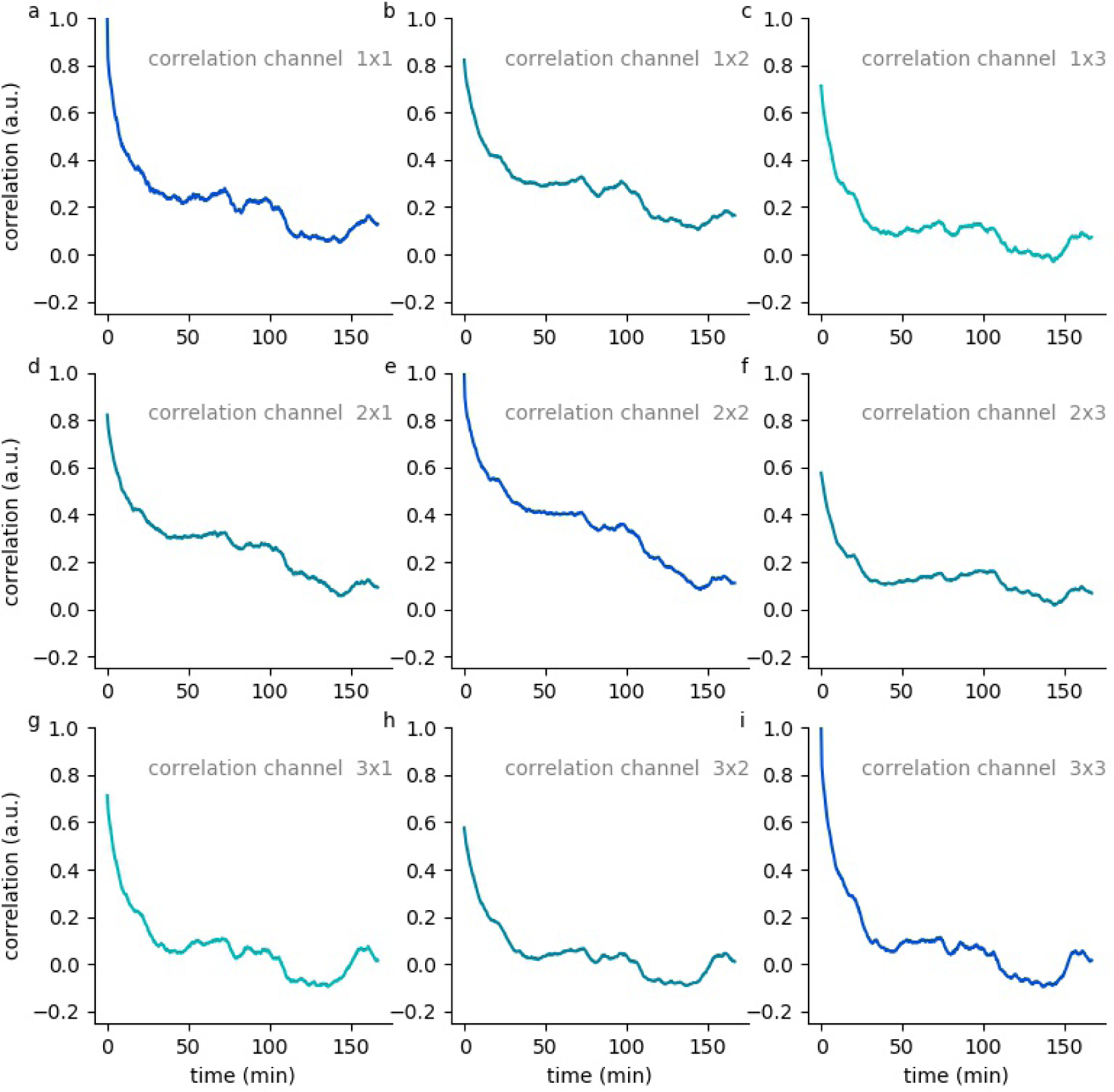
Hypnodensity cycles. Temporal auto- and cross correlations of 5-dimensional hypnodensity probability vectors of sleep stages. Shown are date from subject 63.

**Figure S18:**
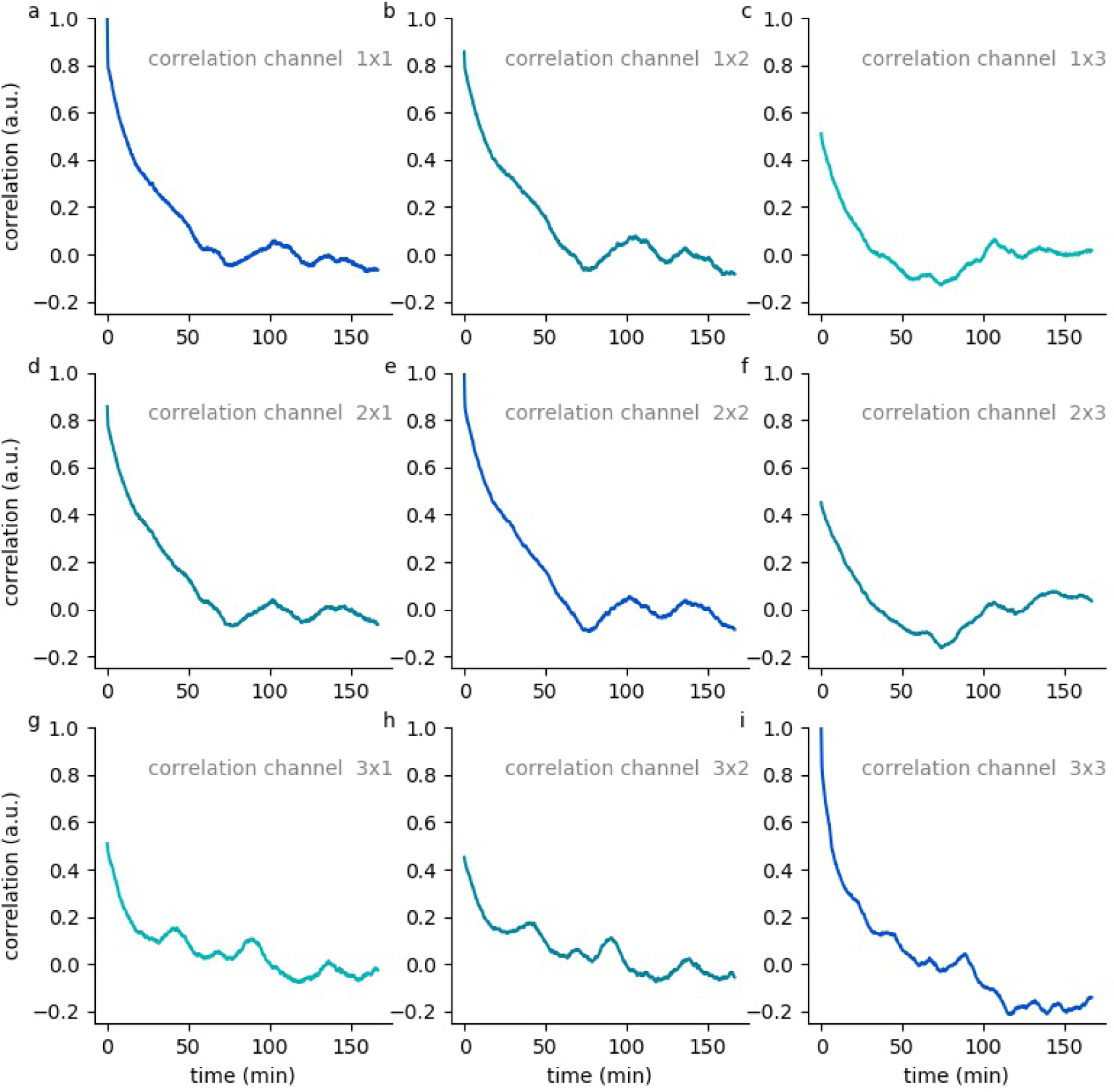
Hypnodensity cycles. Temporal auto- and cross correlations of 5-dimensional hypnodensity probability vectors of sleep stages. Shown are date from subject 64.

**Figure S19:**
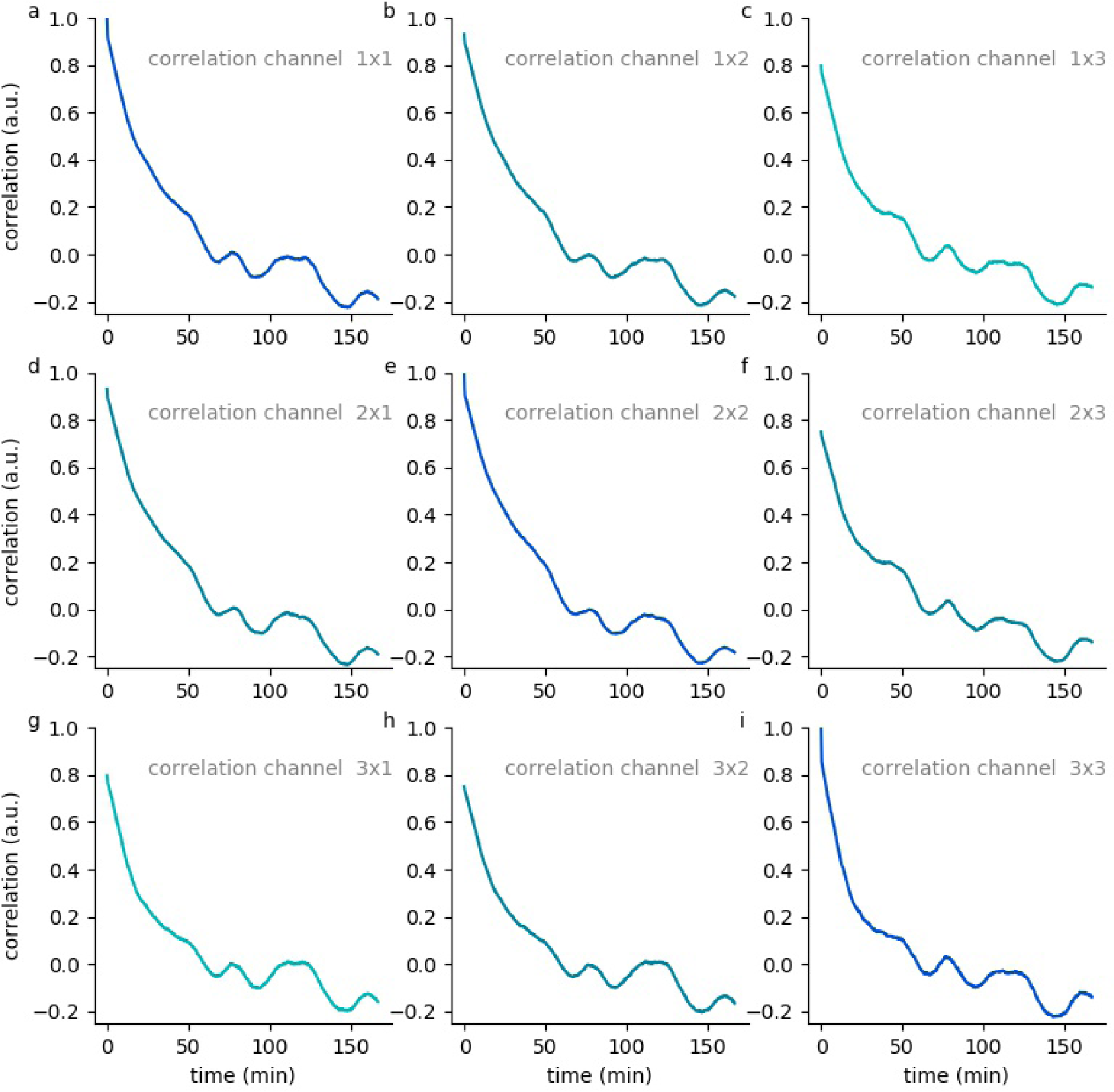
Hypnodensity cycles. Temporal auto- and cross correlations of 5-dimensional hypnodensity probability vectors of sleep stages. Shown are date from subject 65.

